# Influence of solvent, sex, and age on pharmacokinetic and acute behavioral effects of vaporized cannabis extract in mice

**DOI:** 10.1101/2025.01.10.632469

**Authors:** S.R. Westbrook, A.L. Jensen, V. Copeland-Solorzano, J. Buursma, G. Freeby, T. von Melville, T. Edwards, K. Hayashi, R.J. McLaughlin, K.M. Delevich

**Author notes:** **denotes co-first authorship** (S.R. Westbrook, A.L. Jensen). **Corresponding Author** Kristen M. Delevich, Ph.D. Washington State University College of Veterinary Medicine Department of Integrative Physiology and Neuroscience Pullman, WA 99164.

## Abstract

The legalization of cannabis in several states across the US has increased the need to better understand its effects on the body, brain, and behavior, particularly in different populations. Rodent models are particularly valuable in this respect because they provide precise control over external variables. Previous rodent studies have found age and sex differences in response to injected Δ^9^-tetrahydrocannabinol (THC), the major psychoactive component of cannabis. However, this route of administration does not mimic the most common way humans consume cannabis, i.e. through inhalation. Here, we sought to address this gap by investigating age and sex differences in pharmacokinetics and the acute behavioral effects of vaporized cannabis extract in mice. Adolescent (postnatal day [P] 35-50) and adult (≥ P70) mice of both sexes received noncontingent exposure to 0 mg/ml, 150 mg/ml, or 300 mg/ml vaporized cannabis extract diluted in either 80% propylene glycol/20% vegetable glycerol (PG/VG) or 100% polyethylene glycol 400 (PEG). Immediately after exposure, body temperature, hot plate withdrawal latency, and locomotion were assessed. Blood was collected at 0, 30, and 60 min after vapor exposure, and plasma THC and its metabolites were analyzed. Measured THC levels were higher in both the plasma of vapor-exposed mice and the cannabis extract solutions themselves when PEG was the solvent compared to PG/VG. Vaporized cannabis (dissolved in PEG) at the highest dose tested induced hypothermic, antinociceptive, and locomotor-suppressing effects in all groups of mice. We found a dose-dependent age difference in locomotion, indicating that adolescents were less sensitive to the locomotor-suppressing effects of vaporized cannabis, which may be related to the plasma THC levels achieved. Although we found no significant sex differences in the acute behavioral effects of vaporized cannabis, there were significant sex differences in plasma THC metabolites indicating that female mice may metabolize vaporized cannabis more slowly than male mice. Taken together, the current findings add to a growing number of studies implementing vaporized cannabinoid delivery approaches by revealing PEG as the superior solvent for studies involving cannabis extract.

## Introduction

The expansion of recreational cannabis legalization in the United States has been met with decreasing harm perceptions, broader social acceptance, and rising daily cannabis use (Compton et al., 2019). Inhalation is one of the most common routes of cannabis use in humans (Wadsworth et al., 2022; Streck et al., 2019), and cannabis vaping dramatically increased in adolescents from 2017-2019 (Hammond et al., 2021; Wadsworth et al., 2022) and has remained high since (Substance Abuse and Mental Health Services Administration, 2023). Despite the decreasing stigma and rise of cannabis vaping, the pharmacological effects of acute cannabis intoxication from vaping, particularly in different populations, remain poorly understood.

Cannabis from the *Cannabis sativa* plant is composed of over 120 phytocannabinoids (Morales et al., 2017), including THC and cannabidiol (CBD). Human laboratory studies employing vaporized cannabis have shown sex differences in its acute pharmacological effects, with females exhibiting higher plasma levels of THC metabolites and reporting greater sensitivity to subjective drug effects (Sholler et al., 2021). Historically, preclinical studies of cannabinoids have used either synthetic cannabinoids or isolated constituents (i.e., THC) and predominantly administered them via intraperitoneal injections. Notably, pharmacokinetics (Baglot et al., 2021; Manwell et al., 2014a) and behavioral effects (Manwell et al., 2014a,b) greatly differ between injected and inhaled THC. More recently, the vaporized cannabinoid model in rats has gained popularity with multiple studies recapitulating the sex difference in pharmacokinetics reported in humans who use vaporized cannabis—namely, higher 11-hydroxy-Δ^9^-tetrahydrocannabinol (11-OH-THC) plasma levels in females (Lightfoot et al., 2024; Freels et al., 2024; Baglot et al., 2021; Ruiz et al., 2021; Glodosky et al., 2020). In rodents, cannabinoids tend to produce a canonical tetrad of behavioral effects—hypothermia, antinociception, suppressed locomotion, and catalepsy (Little et al., 1988; Metna-Laurent et al., 2017). In general, female rodents tend to exhibit greater behavioral sensitivity to cannabinoid injections compared to males (Craft et al., 2013). However, recent reports using vaporized THC have found no sex differences in vaporized THC-induced hypothermia in adolescent (Ruiz et al., 2021; Nguyen et al., 2020) or adult rats (Baglot et al., 2021), suggesting that sex differences in some tetrad effects may depend on route of administration and plasma THC levels achieved.

Considerably less is known about age differences in the pharmacology of vaporized cannabis, with no studies to date including a direct comparison of adolescents vs. adults. However, one human study reported that young adults (aged 21-25 years old) experienced greater craving and deleterious cognitive effects after smoking THC-dominant flower compared to older adults (aged 55-70), despite achieving similar plasma THC levels (Mueller et al. 2021). One preclinical study reported higher plasma THC metabolites (11-OH-THC and 11-nor-9-carboxy-THC [THC-COOH]) in adolescent relative to adult mice following intraperitoneal injection of THC, although this study only included males (Torrens et al., 2020). Therefore, while existing data suggest that cannabis metabolism and its behavioral effects may differ by age and sex, this has not been systematically assessed using a preclinical model that mimics the composition and route of administration most often used by humans. Furthermore, it will be important to examine potential age and sex effects on cannabis pharmacokinetics and behavioral endpoints, as sex differences in cannabis use emerge during the adolescent to adult transition in humans (Bhatia et al., 2023; Windle, 2020).

We sought to fill this gap in knowledge by using a model of vaporized cannabis exposure (Freels et al., 2024; Weimar et al., 2023; Freels et al., 2020; Glodosky et al., 2020; Weimar et al, 2020) in mice to investigate age and sex differences in the pharmacology of acute cannabis exposure. Additionally, since there is a lack of consensus regarding the optimal solvent to use in vapor delivery studies, we explored behavioral and pharmacological effects of vaporized cannabis extract in two commonly used solvents. Importantly, our model uses a whole-plant derived cannabis extract, instead of isolated THC, which increases the translational relevance by more closely resembling the commercially available broad-spectrum products used by humans. Establishing this model in mice is an important first step to pave the way for future mechanistic investigations that leverage the extensive genetic toolkit available in mice compared to rats. We acutely exposed mice to vaporized vehicle or cannabis extract that differed by dose and solvent and measured the acute behavioral effects and pharmacokinetics via plasma THC and metabolites. We hypothesized that acute exposure to whole-plant cannabis vapor would produce hypothermia, antinociception, and suppressed locomotion in a dose-dependent manner. Importantly, we hypothesized that these effects would be influenced by age and sex with adolescents and females showing greater pharmacological effects of acute vaporized cannabis exposure.

## Methods

### Husbandry

Male and female C57BL/6 (C57) mice obtained from Jackson Laboratories were bred in-house in a climate-controlled vivarium. Offspring used for the experiments were weaned on postnatal day (P) 21 and housed in groups of 2-5 same-sex siblings on a 12:12 h normal light/dark cycle (lights on at 7 h) in standard ventilated polycarbonate cages with Biofresh cellulose bedding. Mice were provided *ad libitum* access to standard chow (Purina 5001) and water in the home cage.

### Experiment

#### Subjects

Experiments were conducted during the light cycle (7 h - 13 h). 414 mice (205 females, 209 males) were run in the experiment with the following design: 2 sex x 2 age x 3 dose x 2 solvent with blood collected at three timepoints (Figure 1). Adolescent (postnatal day [P] 35-50) and adult (≥ P70) mice of both sexes underwent noncontingent vapor exposure to 0 mg/ml, 150 mg/ml, or 300 mg/ml cannabis extract diluted in 80% propylene glycol/20% vegetable glycerol (PG/VG) or 100% polyethylene glycol 400 (PEG) with blood collected at 0-, 30-, or 60-min post-exposure. Since mice were euthanized for blood collection, mice for the 0-min timepoint did not undergo behavioral testing. The 30- and 60-min groups underwent behavioral testing immediately following exposure as described below. All procedures were approved by the Washington State University Institutional Animal Care and Use Committee.

**Figure 1.**
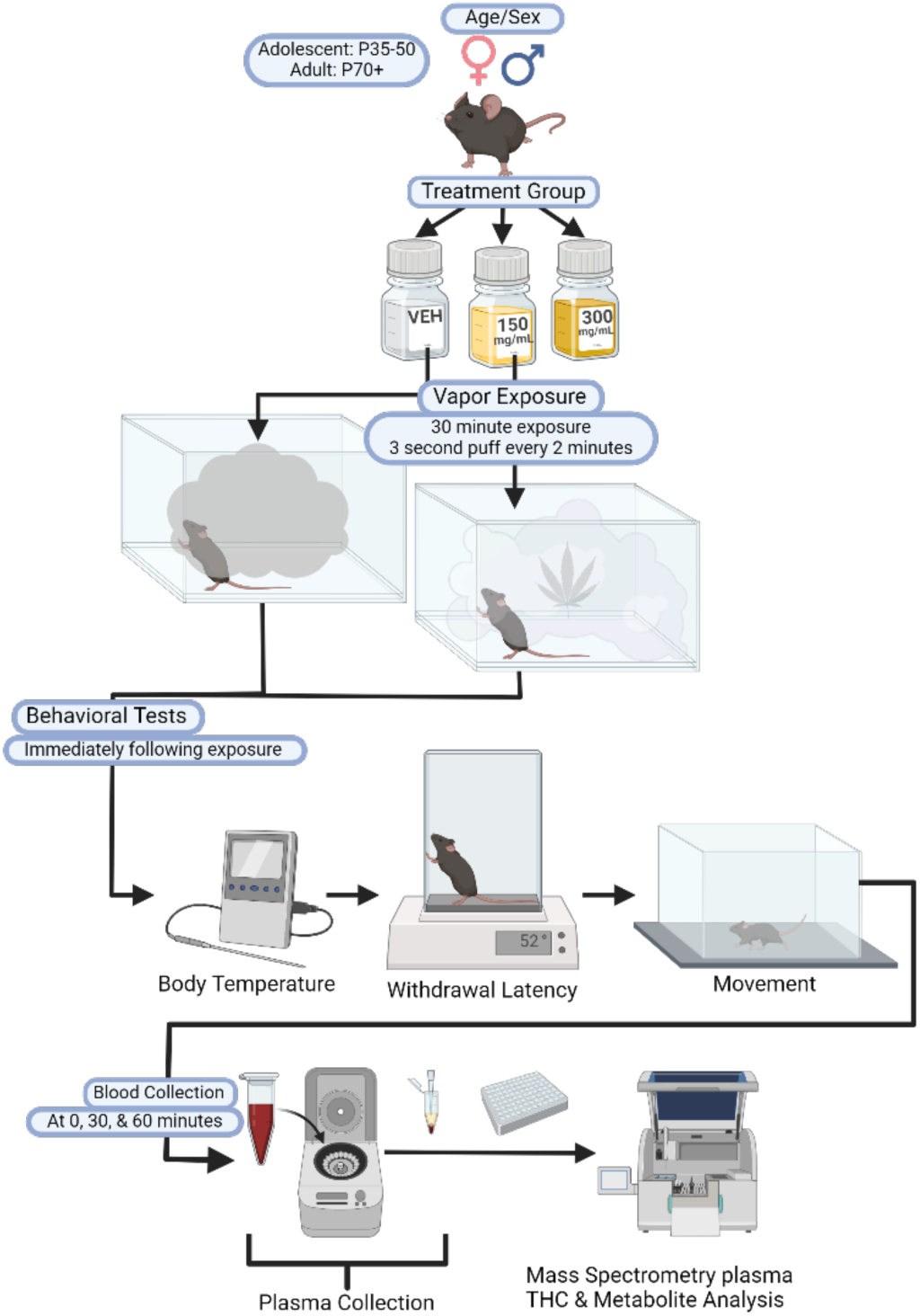
Schematic of the experimental paradigm. Adolescent and adult mice of both sexes were exposed to either vehicle, 150 mg/ml, or 300 mg/ml cannabis vapor for one 30-minute session. Blood was collected either immediately after the session (0-min timepoint) or after completion of behavioral testing (body temperature, hot plate, and open field) at 30 or 60 min after the vapor session. Blood samples were centrifuged, and plasma collected for THC and metabolite measurements via mass spectrometry.

#### Drugs

Whole plant cannabis extract was obtained from the National Institute on Drug Abuse (NIDA) Drug Supply Program. Raw extract contained 60.31% THC, 0.15% CBD, 2.38% cannabigerol (CBG), 2.03% cannabinol (CBN), 1.18% cannabichromene (CBC), 0.78% tetrahydrocannabivarin (THCV), and 0.25% Δ^8^-tetrahydrocannabinol (Δ^8^-THC). Extract was prepared at 150 mg/ml and 300 mg/ml concentrations by dissolving raw cannabis extract into a vehicle (VEH) of 80% propylene glycol/20% vegetable glycerol (PG/VG) or 100% polyethylene glycol 400 (PEG) under continuous stirring at 60^°^ C as described previously (Freels et al., 2020).

#### Experimental Design

##### Pre-Exposure Baselines

On the first experimental day, mice were habituated to the testing room for one hour prior to hot plate exposure (Hot plate Analgesia Meter – Columbus Instruments). Mice were placed within an acrylic cylinder (12.7 cm diameter x 25.4 cm height) on the hot plate, which was heated to 52°C. Hot plate withdrawal latency was determined by the number of seconds it took for the mouse to flick or lick a hind paw or jump. Mice were removed from the hot plate if no response was exhibited within 30 seconds. Pre-exposure hot plate measurements included three trials with an intertrial interval of five minutes. The first pre-exposure trial was used to habituate the mouse to the hot plate, whereas trials two and three were averaged to generate the pre-exposure withdrawal latency.

On exposure day (24-48 hours later), mice were weighed and then habituated to the room for 30 minutes. Body temperatures were taken immediately before and after vapor exposure using a rectal thermometer (Physiotemp digital thermometer) lubricated with Vaseline and inserted 2 cm.

##### Vaporized Cannabis Exposure

Eight 22 x 20 x 14-cm (L x W x H) vapor chambers from La Jolla Alcohol Research Inc. ([LJARI] La Jolla, CA) were controlled by a tablet with custom designed software from LJARI, INC to deliver noncontingent vapor puffs. Air entered through the front of the chambers through a port, and air was removed by a vacuum in the back of the chamber (filtered through Whatman HEPA-CAP filter) in a unidirectional flow. Tubing from the air intake port connected to the airflow meter that was set at 1L/min, which connected to a commercial e-cigarette cartridge (Z Sub-OHM tank 5mL capacity by GEEK VAPE with 0.2 Ω M2 atomizer). Each chamber was equipped with two nose ports and cue lights located on the left and right chamber walls, although the ports and cue lights were not active during the vapor exposure.

One to two mice (cagemates only) were placed in a vapor chamber with an iso-PAD (Braintree Scientific). A three-second vapor puff was administered every two minutes with simultaneous illumination of a cue light. A total of 15 puffs of vapor was administered during the 30-min session.

##### Post-Exposure Behavior

At the end of exposure, 30 and 60-minute blood collection groups had temperatures taken, one trial of post-exposure hot plate conducted, and were put into the open field locomotion chamber, a clear acrylic box (28cm x 28cm x 20.5 cm) (W x L x H). Locomotion was video recorded using a webcam (Logitech C920e HD 1080p) for fifteen minutes.

##### Blood Collection and Metabolites

Mice with 0-minute time points were euthanized for blood collection immediately after vapor exposure. Mice from the 30 and 60 min timepoint groups underwent behavioral testing immediately post-exposure before being euthanized for blood collection at their respective times post-exposure. Mice were euthanized using CO_2_, after which intra-cardiac blood collection was performed. Blood was collected within fifteen minutes of target collection time and put in an Eppendorf tube with 30 µL of EDTA. Samples were centrifuged at 2000 rcf for 15 minutes at 4°C, and plasma was extracted into 100 ul duplicates and stored at -80 °C. Cannabinoids in plasma (THC, 11-OH-THC, and THC-COOH) were analyzed by the Washington State University Tissue Imaging, Metabolomics, and Proteomics Laboratory, Pullman, WA as previously described (Britch et al., 2017) using a Synapt G2-S (Waters) mass spectrometer. Samples with a value of 0 were included, and samples with a value below the limit of detection (<1ng/ml) were excluded from statistical analyses.

#### Data Analysis

All behavioral videos were analyzed for total locomotion (cm) using an offline behavioral tracking software, Deeplab Cut (Mathis et al., 2018) and SimBa (Goodwin et al. 2024). Statistical analyses were performed in SAS OnDemand for Academics: Studio statistical software using proc mixed for analyses of variance, proc reg for linear regressions, and proc freq for chi square tests of independence. Graphs were generated using GraphPad Prism software (version 10, San Diego, CA, USA).

Dependent measures were hot plate withdrawal latency difference score (calculated as post-exposure – pre-exposure average), body temperature difference score (calculated as post-exposure – pre-exposure), and total locomotion (cm) in the open field. For hot plate data, a single trial was excluded if a mouse urinated. If trial 2 or 3 was excluded, the remaining trial was used as the pre-exposure withdrawal latency in the difference score calculation (n=48). If either (1) both trials 2 and 3 were excluded or (2) the single post-exposure trial was excluded, then the hot plate difference score could not be calculated, so all of the hot plate data from that animal was excluded (n=28). The hot plate data from two more mice were excluded due to experimenter error. For body temperature, if either the pre-exposure or post-exposure value was missing, the difference score could not be calculated, so the animal’s body temperature data was excluded (n=4). Total locomotion data for 23 mice were lost due to equipment error. Three mice were excluded from all statistical analyses due to vapor equipment failure. Group information for mice that met exclusion criteria is provided in the Supplemental Table S1. Final sample sizes for behavior and plasma data are provided in Tables 1 and 2, respectively.

**Table 1.** Final sample sizes per group for the three behavioral endpoints.

| Body Temperature | Solvent | Dose | Adolescent Females | Adolescent Males | Adult Females | Adult Males |
| --- | --- | --- | --- | --- | --- | --- |
|  | PG/VG | Vehicle | 11 | 8 | 14 | 15 |
|  |  | CAN150 | 11 | 14 | 20 | 20 |
|  |  | CAN300 | 12 | 14 | 15 | 17 |
|  |  |  |  |  | Total n: 171 |  |
|  | PEG | Vehicle | 13 | 12 | 11 | 12 |
|  |  | CAN150 | 12 | 12 | 12 | 13 |
|  |  | CAN300 | 12 | 12 | 12 | 13 |
|  |  |  |  | Total n: 146 |  |  |
| Hot Plate | PG/VG | Vehicle | 11 | 7 | 13 | 11 |
|  |  | CAN150 | 10 | 12 | 16 | 19 |
|  |  | CAN300 | 11 | 11 | 14 | 14 |
|  |  |  |  |  | Total n: 149 |  |
|  | PEG | Vehicle | 13 | 12 | 11 | 12 |
|  |  | CAN150 | 10 | 12 | 15 | 12 |
|  |  | CAN300 | 11 | 12 | 12 | 11 |
|  |  |  |  |  | Total n: 143 |  |
| Locomotion | PG/VG | Vehicle | 9 | 8 | 14 | 15 |
|  |  | CAN150 | 9 | 14 | 17 | 14 |
|  |  | CAN300 | 12 | 14 | 13 | 15 |
|  |  |  |  |  | Total n: 154 |  |
|  | PEG | Vehicle | 13 | 12 | 11 | 12 |
|  |  | CAN150 | 12 | 12 | 10 | 13 |
|  |  | CAN300 | 12 | 12 | 12 | 13 |
|  |  |  |  |  | Total n: 144 |  |

**Table 2.**
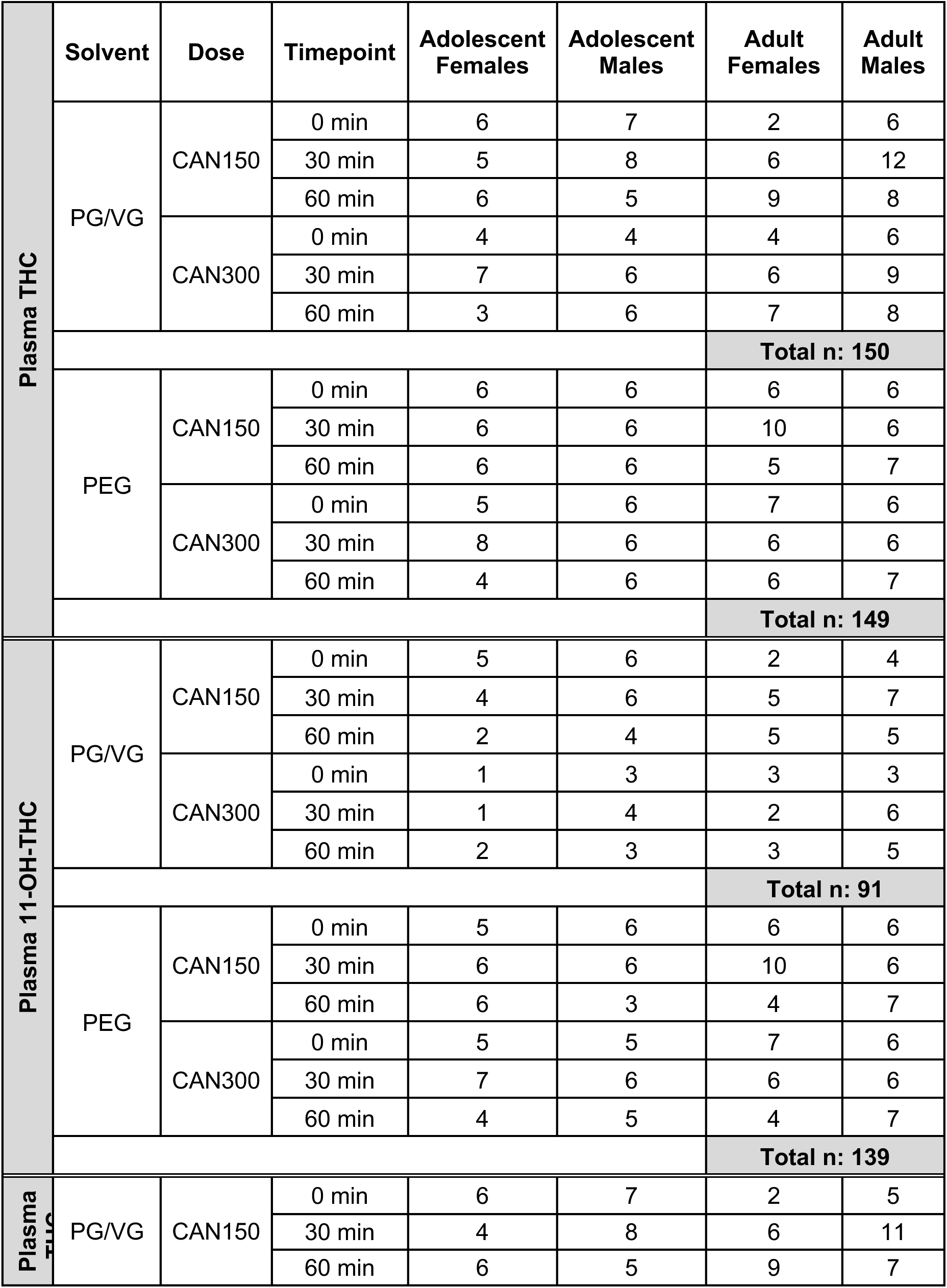
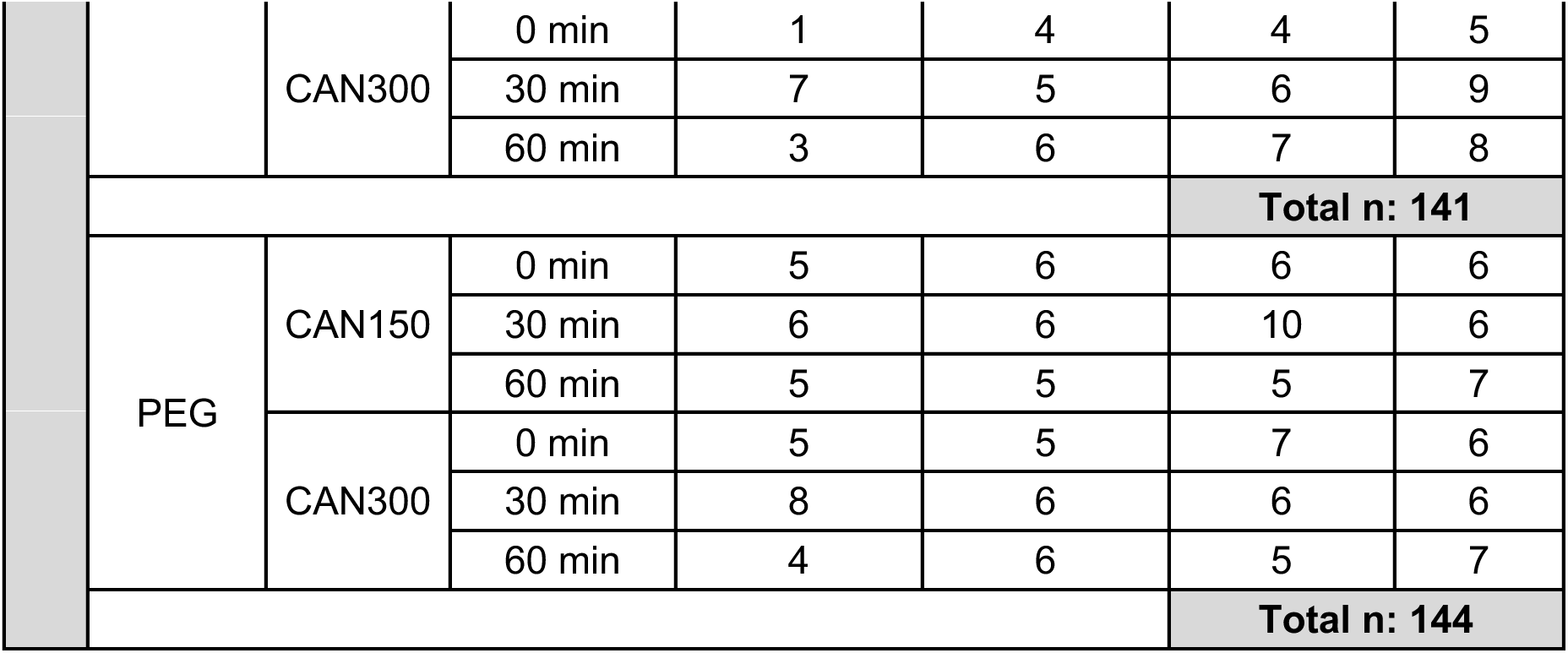
Final sample sizes per group for plasma THC and its metabolites, 11-OH-THC and THC-COOH.

All factors (Sex, Age, Dose, Solvent, Timepoint) were between subjects. Chi-square tests of independence were conducted on plasma data to determine whether there was an association between Solvent and the proportion of samples with values below the level of detection (<1 ng/ml). Five-way ANOVAs were conducted on plasma levels of THC, 11-OH-THC, and THC-COOH with Sex, Age, Dose (150 mg/ml, 300 mg/ml), Solvent, and Timepoint (0 min, 30 min, 60 min) as factors. Four-way analyses of variance (ANOVAs) were used to assess the interaction of Sex, Age, Dose (0 mg/ml, 150 mg/ml, 300 mg/ml), and Solvent (PG/VG, PEG) on body temperature difference score, hot plate difference score, and total locomotion. When significant effects of Solvent were detected, separate analyses were conducted for each solvent. Z-scores for plasma THC levels were calculated within each timepoint (i.e., 30 min or 60 min) for each solvent, so that timepoints could be collapsed for separate (by solvent) linear regression analyses assessing plasma THC levels (z-scores) as a predictor for each behavioral endpoint. P-values below 0.05 were considered statistically significant, and significant effects were followed up with Tukey’s post-hoc comparisons where appropriate.

## Results

### Plasma THC

Blood was collected at three timepoints after acute cannabis vapor (CAN 150, CAN 300) exposure (0 min, 30 min, or 60 min) for analyses of plasma levels of THC and its metabolites, 11-OH-THC and THC-COOH. Chi Square tests of independence revealed that plasma levels below the level of detection (<1 ng/ml) were significantly more common when the solvent was PG/VG compared to PEG [Supplementary Tables S2-S4; Plasma THC: *Χ*^2^(1, *N*=314) = 10.6670, *p*=0.0011; Plasma 11-OH-THC: *Χ*^2^(1, *N*=314) = 55.2650, *p*<0.0001; Plasma THC-COOH: *Χ*^2^(1, *N*=314) = 9.3916, *p*=0.0022]. Plasma levels below detection were excluded from the remaining statistical analyses.

A 5-way ANOVA (Age, Sex, Dose, Timepoint, and Solvent) on plasma THC levels revealed a significant 5-way interaction [F_2,251_ = 4.83, *p*=0.0088; Supplemental Table S5], and therefore Tukey’s post-hoc comparisons for significant simple main effects were examined (Supplemental Table S6). Expected significant effects of Timepoint, whereby the 0 min timepoint had the highest plasma THC levels followed by 30 min and 60 min timepoints, were present in all Age x Sex x Dose groups with PEG as the solvent. Notably, there were no significant effects of Timepoint for mice with PG/VG as the solvent. At the 0 min timepoint, plasma THC levels were significantly higher when the solvent was PEG compared to PG/VG for almost all Age x Sex x Dose groups.

The greater proportion of plasma samples below the level of detection and the lack of robust effects of Timepoint on plasma THC levels when PG/VG was the solvent led us to test how the solvent used affected THC levels measured in the prepared cannabis extract solution. We compared THC levels immediately after cannabis extract was dissolved in PG/VG vs. PEG and across time to determine whether there were differences in how readily the cannabis extract went into and stayed in solution. We performed a 3-way mixed ANOVA with Dose (150 mg/ml, 300 mg/ml) and Solvent (PG/VG, PEG) as between subjects factors and Day (0, 3, 14) as the within subjects factor. The THC levels in solution were significantly higher when cannabis extract was dissolved in PEG compared to PG/VG (Supplemental Figure S1), as indicated by a main effect of Solvent [F_1,2_ = 26.25, *p*=0.0360]. Since (1) the *in vivo* plasma THC and metabolites levels were more often below the level of detection with PG/VG compared to PEG, (2) the plasma THC levels from mice with PG/VG as the solvent lacked expected robust Timepoint effects, (3) the solution levels of THC were significantly lower in cannabis extract dissolved in PG/VG compared to PEG, and (4) we found consistent main effects of Solvent for plasma THC and all behavioral measures (Supplemental Tables S5-S9), we chose to focus on the PEG analyses for the remainder of this study, but provide the PG/VG data analysis in the Supplemental Materials.

Plasma THC levels for mice with PEG as the solvent are shown in Figure 2A, B. Analysis of plasma THC levels with a 4-way ANOVA (Age, Sex, Dose, and Timepoint) revealed significant Age x Dose [F_1,125_ = 4.96, *p*=0.0277], Age x Sex x Timepoint [F_2,125_ = 7.44, *p*=0.0009], and Sex x Dose x Timepoint [F_2,125_ = 3.37, *p*=0.0376] interactions. Simple main effects for the Age x Dose interaction indicate expected dose-dependent increases in plasma THC levels for adolescents (post-hoc: CAN150 < CAN300, *p*<0.0001) and adults (post-hoc: CAN150 < CAN300, *p*=0.0379). Further analysis of the Age x Sex x Timepoint interaction showed that at the 0 min timepoint adult males had higher plasma THC levels than adolescent males (post-hoc: *p*=0.0058). As expected, within each Age x Sex group, there was a significant effect of Timepoint, whereby plasma THC levels peak at 0 min with no significant differences between 30 min and 60 min (post-hocs within each Age x Sex group: 0 min > 30 min, all *p*’s<0.0001, 0 min > 60 min, all *p*’s<0.0001). Further analysis of the Sex x Dose x Timepoint interaction indicated expected dose-dependent increases in plasma THC levels for both male (post-hoc: CAN150 < CAN300, *p*=0.0063) and female mice (post-hoc: CAN150 < CAN300, *p*<0.0001) at the 0 min timepoint. Finally, within each Sex x Dose group, there was an expected effect of Timepoint, whereby plasma THC levels were the highest at 0 min with no significant differences between plasma THC levels at 30 min and 60 min (post-hocs within each Sex x Dose group: 0 min > 30 min, all *p*’s<0.0001, 0 min > 60 min, all *p*’s <0.0001).

**Figure 2.**
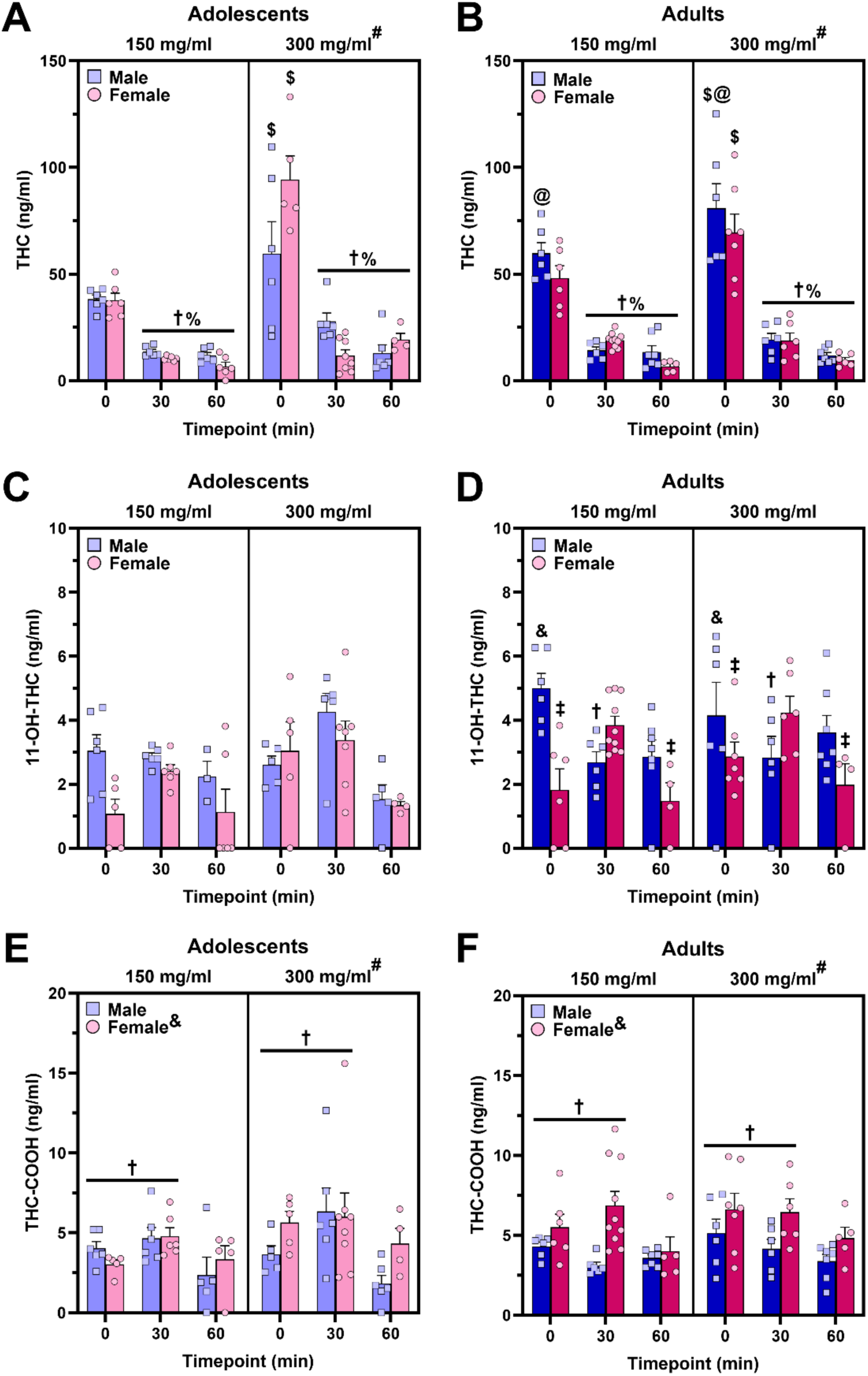
Time course of plasma THC and metabolites following 30-min vapor exposure session. (**A,B**) Plasma THC levels showed expected dose-dependent increases (*post-hoc*: ^#^*p*<0.05, vs. CAN150 within age; ^$^*p*<0.01, vs. CAN150 within sex and timepoint collapsed across age) and time-dependent decreases (*post-hoc*: ^†^*p*<0.001, vs. 0 min timepoint within age and sex collapsed across dose; ^%^*p*<0.001, vs. 0 min timepoint within sex and dose collapsed across age). Adult male mice, regardless of dose, reached higher plasma THC levels at the 0 min timepoint than their adolescent counterparts (*post-hoc*: ^@^*p*<0.01, vs. adolescent males within timepoint collapsed across dose). (**C,D**) Adult male mice reached a significantly higher and earlier peak in plasma 11-OH-THC levels at the 0 min timepoint (*post-hoc*: ^&^*p*<0.01, vs. adult females at the 0 min timepoint collapsed across dose; ^†^*p*<0.05, vs. 0 min timepoint within adult males collapsed across dose) compared to females who reached their peak at the 30 min timepoint (*post-hoc*: ^‡^*p*<0.05, vs. 30 min timepoint within adult females collapsed across dose). No significant sex differences were seen for plasma 11-OH-THC levels in adolescent mice. (**E,F**) Females reached higher plasma THC-COOH levels compared to males (main effect of Sex: ^&^*p*<0.001). Mice exposed to 300mg/ml cannabis vapor had higher levels of THC-COOH compared to those exposed to the lower dose (main effect of Dose: ^#^*p*<0.05). A significant main effect of Timepoint indicated that plasma levels of THC-COOH significantly dropped off at 60-min relative to the 0-min and 30-min timepoints (*post-hoc*: ^†^*p*<0.05, vs. 60 min timepoint).

### Plasma THC Metabolites

Separate 4-way ANOVAs were conducted on levels of plasma THC metabolites—namely, 11-OH-THC and THC-COOH—with Age, Sex, Dose, and Timepoint as between subjects factors. Analysis of plasma levels of 11-OH-THC (Figure 2C, D) revealed a significant Age x Sex x Timepoint interaction [F_2,115_ = 5.71, *p*=0.0043]. Further analyses indicate that 11-OH-THC plasma levels significantly peak at 30 min for adult females (post-hoc: 0 min < 30 min, *p*=0.0447; 30 min > 60 min, *p*=0.0063) and at 0 min for adult males (post-hoc: 0 min > 30 min, *p*=0.0447). At the 0 min timepoint, adult males achieved significantly higher plasma 11-OH-THC levels compared to adult females (post-hoc: *p*=0.0029). By contrast, significant sex differences were not seen within the adolescent group. In general, these adult sex differences reflect that adult males reach their peak plasma 11-OH-THC earlier than adult females, suggesting that female mice metabolize vaporized cannabis more slowly than male mice in adulthood.

Plasma levels of THC-COOH (Figure 2E, F) showed dose-dependent increases [main effect of Dose: F_1,120_ = 4.41, *p*=0.0377]. Female mice had higher THC-COOH levels compared to male mice [main effect of Sex: F_1,120_ = 12.55, *p*=0.0006]. A main effect of Timepoint [F_2,120_ = 9.56, *p*=0.0001] indicated that plasma levels of THC-COOH drop off at 60 min as the levels at 0 min [post-hoc: *p*=0.0118] and 30 min [post-hoc: *p*<0.0001] were significantly higher than the levels at 60 min with no significant differences between 0 min and 30 min timepoints.

### Behavior

Linear regression analyses of plasma THC levels (z-scores) for mice with PEG as the solvent indicated that plasma THC was significantly correlated with all three behavioral outcomes—body temperature difference [adj r^2^=0.1302, *p*=0.0002; Figure 3A], hot plate withdrawal latency difference [adj r^2^=0.0879, *p*=0.0022; Figure 3C], and total movement [adj r^2^=0.1166, *p*=0.0004; Figure 3E]. Higher plasma THC levels predicted lower body temperature difference scores (i.e., more hypothermia), higher hot plate difference scores (i.e., more antinociception), and lower total movement in the open field (i.e., more hypolocomotion). Moreover, all three behavioral measures were significantly correlated with each other (Table 3).

**Figure 3.**
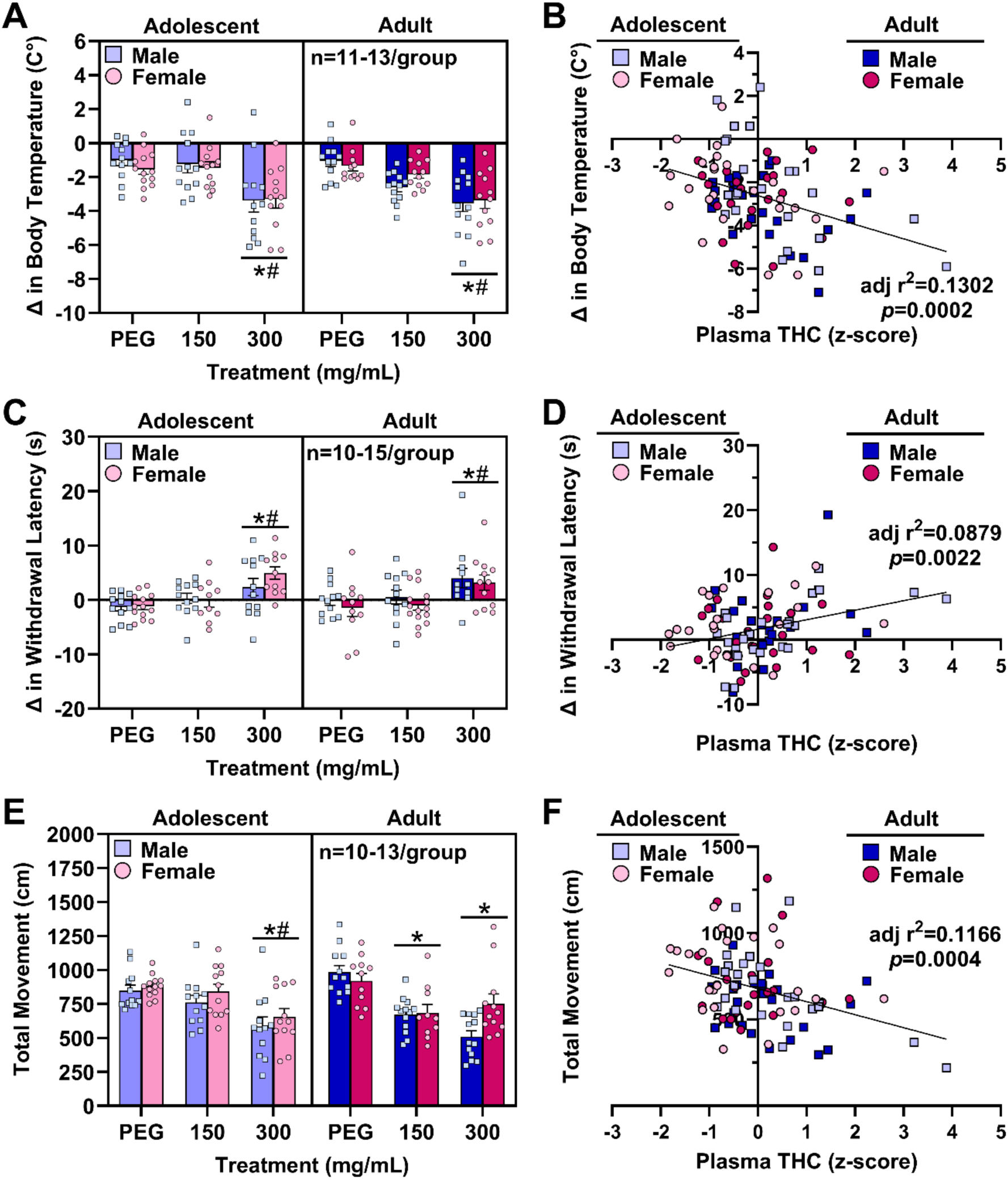
Vaporized cannabis extract dissolved in PEG produced dose-dependent hypothermia, antinociception, and hypolocomotion in adolescent and adult mice of both sexes, and these effects were significantly correlated with plasma THC levels. (**A**) Mice exposed to 300 mg/ml cannabis vapor exhibited greater hypothermia relative to mice exposed to 150 mg/ml cannabis vapor (*post-hoc:* ^#^*p*<0.001) or vehicle vapor (*post-hoc: *p*<0.001), regardless of age or sex. (**B**) Plasma THC levels significantly predicted changes in body temperature. (**C**) Mice exposed to 300 mg/ml cannabis vapor displayed greater antinociception relative to mice exposed to the lower dose of cannabis vapor (*post-hoc:* ^#^*p*<0.001) or vehicle vapor (*post-hoc: *p*<0.001), regardless of age or sex. (**D**) Plasma THC levels significantly predicted changes in withdrawal latency in the hot plate test. (**E**) Adult mice exposed to either dose of cannabis vapor moved significantly less compared to their vehicle-exposed counterparts (*post-hoc:* \**p*<0.001 vs. vehicle). Only adolescent mice exposed to the higher dose of cannabis vapor exhibited significant hypolocomotion (*post-hoc:* \**p*<0.001 vs. vehicle; ^#^*p*<0.001 vs. CAN150). (**F**) Plasma THC levels significantly predicted locomotor activity.

**Table 3.** Adjusted r^2^ and *p*-values for linear regressions between the three behavioral endpoints, which indicate that each measure significantly predicted the other measures.

|  |  | **Outcome** |  |  |
| --- | --- | --- | --- | --- |
| | **Adj $r^2$**<br>**(\*$p$-value)** | $\Delta$ in Body Temperature | $\Delta$ in Withdrawal Latency | Total Movement |
| **Predictor** | $\Delta$ in Body Temperature | NA | **0.1894**<br>**(\*<0.0001)** | **0.2198**<br>**(\*<0.0001)** |
| | $\Delta$ in Withdrawal Latency | **0.1894**<br>**(\*<0.0001)** | NA | **0.1251**<br>**(\*0.0004)** |
|  | Total Movement | **0.2198**<br>**(\*<0.0001)** | **0.1251**<br>**(\*0.0004)** | NA |

Group differences in behavioral measures when PEG was the solvent were analyzed with 3-way (age, sex, dose) ANOVAs. Mice exposed to the higher dose of cannabis vapor (CAN300) had more negative body temperature difference scores [main effect of Vapor: F_2,134_ = 28.02, *p*<0.0001] indicating a hypothermic effect relative to CAN150 and vehicle vapor (post-hoc: *p*’s<0.0001; Figure 3B). Exposure to CAN300 vapor also produced a more positive hot plate difference score [main effect of Vapor: F_2,131_ = 16.36, *p*<0.0001], indicating greater antinociception after vapor exposure compared to CAN150 and vehicle vapor-exposed mice (post-hoc: *p*’s<0.0001; Figure 3D). In the open field, female mice moved significantly more than male mice [main effect of Sex: F_1,132_ = 4.27, *p*=0.0407; Figure 3F]. There was a significant Age x Vapor interaction [F_2,134_ = 28.02, *p*<0.0001], reflecting that significant cannabis-induced hypolocomotion was present at both doses for adults (post-hoc: vehicle vs. CAN150 and vehicle vs. CAN300, *p*’s<0.0001), but only at the higher dose for adolescents (post-hoc: vehicle vs. CAN300, *p*<0.0001).

## Discussion

The current study sought to use the whole-plant derived cannabis vapor exposure model in mice to determine whether there are age and sex differences in the acute behavioral effects and pharmacokinetics of vaporized cannabis. We first discovered that PEG was a better solvent for the cannabis extract than PG/VG. Vaporized cannabis dissolved in PEG induced significant hypothermia, antinociception, and hypolocomotion at the highest dose tested (CAN300) with no significant sex differences. The only age difference in behavior we found was dose-dependent, whereby adults displayed vaporized cannabis-induced hypolocomotion regardless of dose, but only the highest dose suppressed locomotion in adolescents. Immediately after vapor exposure, adult males reached higher plasma THC levels compared to adolescent males, which could contribute to the behavioral difference. The current study found sex differences in plasma THC metabolites that suggest females (and particularly adult females) may metabolize vaporized cannabis more slowly than males.

Both PEG and PG/VG are commonly used solvents in vaping products for humans (Traboulsi et al., 2020). In the current study, we found that more of the whole-plant derived cannabis extract went into solution when PEG was the solvent compared to PG/VG. We also observed more variable results in plasma THC and metabolites, as well as acute behavioral effects, with PG/VG relative to PEG as the solvent. Vaporized cannabis with PEG as the solvent produced the canonical cannabinergic behavioral effects of hypothermia, antinociception, and hypolocomotion, while vaporized cannabis in PG/VG did not for most groups. Moreover, levels of plasma THC were higher and THC metabolites were more frequently above the level of detection when PEG was the solvent compared to PG/VG. For all of these reasons, we focus the remainder of the discussion on our findings when PEG was the solvent.

For all behavioral measures, we observed the expected effects characteristic of cannabinoids (i.e., hypothermia, antinociception, hypolocomotion), and all behaviors were predictive of one another, demonstrated by linear regressions. The effects of cannabis vapor exposure were dose-dependent and only evident with CAN300 exposure—the highest dose tested in the current study. One potential limitation of the current study is the assessment of only thermal nociception and not mechanical. While thermal nociception results may be influenced by our observed hypothermic response, many studies have shown cannabinoid-induced antinociception for both mechanical and thermal stimuli (Craft et al., 2019). Another potential limitation is that the typical cannabinoid tetrad assessment includes the bar test to examine cataleptic effects. Previous work in mice has demonstrated that injected THC produced cataleptic effects, while inhaled THC failed to do so (Marshell et al., 2014). Given this previous report and our own observation during a pilot study that vaporized cannabis did not produce catalepsy, we did not include the bar test in the current study.

The current study did not find evidence for significant age or sex differences in cannabis-induced changes in body temperature or antinociception. This lack of age and sex differences contrasts with the previous literature (Torrens et al., 2020; Craft et al., 2019; Wiley et al., 2007), which has reported age and sex differences in the acute effects of injected synthetic cannabinoid agonists or isolated THC. Notably, route of administration (intraperitoneal injection vs. inhalation) has been shown to greatly influence plasma and brain levels of THC and its metabolites in rats (Baglot et al., 2021). Specifically, inhaled THC produces a more rapid peak in plasma and brain THC levels compared to injected THC. Importantly, injected THC compared to inhaled THC leads to higher and more sustained levels of the metabolite, 11-OH-THC, in the plasma and brain, which amplifies the magnitude of the sex difference in this psychoactive metabolite and may lead to more pronounced sex differences in the behavioral effects. Indeed, other studies using aerosolized THC in rats (Ruiz et al., 2021; Javadi-Paydar et al., 2018) have reported no sex differences in THC-induced hypothermia, antinociception, or locomotor suppression consistent with this current report. Thus, the route of administration in the model may explain our lack of sex differences in the acute behavioral effects of vaporized cannabis.

In the current study, we found significant sex differences in THC metabolism following vaporized cannabis exposure in mice; although, these sex differences did not appear to influence the acute behavioral effects as discussed above. Notably, the sexes did not differ in plasma THC levels achieved; rather, sex differences were evident in THC’s two primary metabolites, 11-OH-THC and THC-COOH. Sex effects were most prominent in adult mice, with adult females reaching their peak plasma 11-OH-THC levels later than adult males. Moreover, females had higher levels of plasma THC-COOH than males. These findings are consistent with previous studies in rats using vaporized cannabinoids, where THC metabolite levels were higher in female rats relative to males in adolescence (Ruiz et al., 2021) and adulthood (Lightfoot et al., 2024; Glodosky et al., 2020). The current study in mice demonstrated that immediately after a 30 min exposure session, adult male mice have higher plasma 11-OH-THC compared to their female counterparts that reach their peak levels 30-min later. This finding along with the higher levels of plasma THC-COOH in females suggests that male mice metabolize THC more readily than females.

The current study assessed the acute locomotor-suppressing effects of vaporized cannabis and found significant age differences that were dose-dependent. Notably, adolescent mice exposed to the highest dose of cannabis vapor displayed hypolocomotion compared to their vehicle-exposed counterparts, while adult mice exposed to either dose of cannabis vapor had suppressed locomotion relative to vehicle-exposed adults. This dose-dependent age difference in cannabis-induced hypolocomotion could be explained by age differences in pharmacokinetics and/or pharmacodynamics. In one study (Torrens et al., 2020), liver microsomes from adolescent male mice metabolized THC more readily than those from adult male mice. Here, we found that adolescent male mice reached significantly lower plasma THC levels compared the adult male mice immediately after exposure. Thus, it is possible that the acute locomotor-suppressing effects of vaporized cannabis may subside more quickly in adolescent mice exposed to the low dose if they metabolize THC at a faster rate than adult mice. Alternatively, this age difference may have been most apparent in the locomotor test due to the order of testing. The acute behavioral effects were always assessed in the same order—body temperature, antinociception, and locomotion. We found a dramatic drop in plasma THC levels between 0 min and 30 min post-exposure, and all behavioral measures were completed by 20 min post-exposure. Thus, it is possible that differences in metabolism may have greater effects on behavioral measures taken further from the end of exposure (i.e., locomotion compared to body temperature and the hot plate test). Our study design limited the direct analysis of the role of group differences in plasma THC levels on our behavioral endpoints because (1) mice that underwent behavioral testing were euthanized for blood collection either at 30 min or 60 min post-exposure and (2) plasma THC was not assessed in mice exposed to vehicle vapor. Thus, we did not have plasma THC data closest to the end of the behavioral testing (i.e., 30 min timepoint) for all mice to address the question of the role of plasma THC levels in the acute behavioral effects most appropriately. Rather, we chose to maximize our ability to detect time course differences in the pharmacokinetics of vaporized cannabis. Thus, future work would be needed to confirm whether the differences in plasma THC between adolescents and adults exposed to the low dose mediate the age difference in the acute locomotor-suppressing effects of vaporized cannabis.

Another potential explanation is that adolescents may be less sensitive to the locomotor-suppressing effects of vaporized cannabis due to differences in pharmacodynamics. A previous study similarly found that adolescent male rats were less sensitive to locomotor suppressive effects of injected THC compared to adult rats. In addition, adolescent rats exhibited reduced adrenocorticotropic hormone following THC injection compared to adults, suggesting that the drug experience was less aversive in adolescents compared to adults (Schramm-Sapyta 2007). While we did not assess pharmacodynamics in the current study, we speculate that motor-related brain areas, like the striatum, may be impacted by THC, as well as other phytocannabinoids present in the whole-plant cannabis extract, through their actions at cannabinoid receptors. Cannabinoid receptor 1 (CB1R) levels change across adolescent development in many brain regions (Gomes et al., 2018; Ellgren et al., 2008), and enhanced CB1R function is thought to underlie adolescent-typical behaviors (Schneider et al. 2015). Moreover, dopamine receptors in brain regions involved in motor behaviors change from adolescence to adulthood (Andersen et al., 1997), and CB1R in the striatum negatively modulates dopamine-mediated motor behaviors (Martin et al., 2008). The phytocannabinoids present in our whole plant cannabis extract can stimulate CB1Rs in motor-related brain areas to impact locomotion with age differences in these effects potentially arising from age differences in CB1R expression/function and/or age differences downstream in the dopaminergic system. Future investigations are needed to determine the role of pharmacokinetics and pharmacodynamics in the age difference in vaporized cannabis-induced hypolocomotion. More generally, understanding age-related differences in potential use-limiting effects of cannabis such as hypolocomotion, may help explain escalated cannabis use in adolescents (Mokrysz et al. 2016).

The current study established a model of vaporized cannabis extract in adolescent and adult mice of both sexes and demonstrated that cannabis vapor produces acute hypothermic, antinociceptive, and locomotor-suppressing effects in all groups. We determined that PEG was a better solvent for whole-plant derived cannabis extract relative to PG/VG. Moreover, we found that vaporized cannabis extract is metabolized more slowly in female mice compared to male mice; however, this sex difference did not translate to sex differences in the acute behavioral effects of vaporized cannabis. Finally, adolescent mice were equally or less sensitive to the acute behavioral effects of vaporized cannabis compared to adult mice, depending on the behavior. Overall, the current study adds to a growing number of studies implementing vaporized cannabinoid delivery approaches by revealing PEG as the superior solvent for studies involving cannabis extract and demonstrating the typical acute cannabinoid-induced behavioral effects in mice. By further establishing this model in mice, this will allow future studies to take advantage of the extensive genetic toolkit available in this species for mechanistic investigations.

## Supporting information

Supplemental results

## ACKNOWLEDGEMENTS

The authors would like to thank Maury Cole and LJARI, Inc. for their continued support with vapor chamber troubleshooting and optimization. The authors would also like to thank Qing Wang for her assistance with colony maintenance and Courtney Klappenbach for her assistance with Deep Lab Cut and SimBA methodologies. Finally, the authors would like to thank Dr. Anna Berim and the Washington State University Tissue Imaging, Metabolomics, and Proteomics Laboratory for their excellent technical assistance with analyzing plasma samples.

## AUTHORSHIP CONTRIBUTION STATEMENT

**SRW:** writing-original draft (lead), writing-review & editing (equal), conceptualization (supporting), formal analysis (lead), visualization (lead), supervision (supporting); **ALJ:** writing-original draft (supporting), writing-review & editing (supporting), visualization (supporting), investigation (equal), data curation (equal), supervision (supporting); **VCS:** investigation (equal), data curation (equal); **JB:** investigation (equal), methodology (equal); **GF:** investigation (supporting), methodology (equal); **TvM:** investigation (supporting); **TE:** investigation (supporting); **KH:** conceptualization (equal), funding acquisition (equal); **RJM:** writing-review & editing (equal), conceptualization (equal), funding acquisition (lead), project administration (supporting), supervision (supporting); **KMD:** writing-review & editing (equal), conceptualization (equal), funding acquisition (equal), project administration (lead), supervision (lead)

## CONFLICT OF INTEREST STATEMENT

The authors have no competing personal or financial interests or any other conflicts of interest to disclose.

## FUNDING STATEMENT

This study was supported by an NIH R21 grant from the National Institute on Drug Abuse (R21DA057245-01) awarded to KMD, RJM, and KH, as well as funds provided for medical and biological research by the State of Washington Initiative Measure No. 502 (KMD, RJM, KH). SRW is supported by an NIH F32 fellowship from the National Institute on Drug Abuse (F32DA060685).

## Supplemental Results

### PG/VG as the Solvent

#### Plasma THC

Plasma THC levels for mice with PG/VG as the solvent are shown in Figure S2A, B. Analysis of plasma THC levels with a 4-way ANOVA (Age, Sex, Dose, and Timepoint) revealed significant 4-way interaction [F_2,126_ = 7.53, *p*=0.0008]. Tukey’s post-hoc analyses indicated that adolescent males exposed to 300 mg/ml cannabis vapor reached higher plasma THC levels at 0 min relative to their female counterparts (post-hoc: *p*=0.0068). There were also significant time-dependent decreases in plasma THC levels with the highest level measured at the 0 min timepoint (post-hoc: Adolescent F CAN150: 0 min > 30 min, *p*=0.0051, 0 min > 60 min, *p*=0.0005; Adolescent M CAN150: 0 min > 30 min, *p*<0.0001, 0 min > 60 min, *p*<0.0001; Adolescent M CAN300: 0 min > 30 min, *p*=0.0190, 0 min > 60 min, *p*=0.0011; Adult F CAN300: 0 min > 30 min, *p*=0.0600; 0 min > 60 min, *p*=0.0029; Adult M CAN150: 0 min > 30 min, *p*<0.0001, 0 min > 60 min, *p*<0.0001). It is important to note that in contrast to plasma THC levels when PEG was the solvent, these timepoint effects were not present in all groups for each dose when PG/VG was the solvent. Moreover, at the 0 min timepoint, plasma THC levels were higher in adolescent female (post-hoc: *p*=0.0141) and adult male mice (post-hoc: *p*=0.0012) exposed to 150 mg/ml cannabis vapor compared to 300 mg/ml cannabis vapor—a dose-dependent effect in the opposite direction as was expected. These findings may provide further examples of the instability issue of cannabis extract when dissolved in PG/VG.

The large proportion of samples with plasma THC metabolites—11-OH-THC and THC-COOH—with values below the limit of detection when the solvent was PG/VG precluded our analysis of plasma THC metabolites.

#### Behavior

Linear regression analyses of plasma THC levels (z-scores) for mice with PG/VG as the solvent indicated that plasma THC was not significantly correlated with any of the three behavioral outcomes—body temperature difference [adj r^2^=-0.0087, *p*=0.8185; Supplemental Figure S3A], hot plate withdrawal latency difference [adj r^2^=-0.0083, *p*=0.6388; Supplemental Figure S3C], and total movement [adj r^2^=-0.0029, *p*=0.3973; Supplemental Figure S3E].

Group differences in behavioral measures when PG/VG was the solvent were analyzed with 3-way (age, sex, dose) ANOVAs. A significant Sex x Vapor interaction [F_2,159_ = 3.29, *p*=0.0398] for body temperature difference scores reflected that male mice exposed to either dose of cannabis vapor had lower body temperature difference scores compared to mice exposed to vehicle vapor, indicating a lower body temperature post-exposure (post-hoc: CAN300 vs. vehicle, *p*=0.0011, CAN150 vs. vehicle, *p*=0.0192; Supplemental Figure S3B). Males had higher hot plate difference scores compared to females [main effect of Sex: F_1,137_ = 5.73, *p*=0.0180]. Exposure to CAN300 vapor produced a higher hot plate difference score in adolescent mice [Age x Vapor interaction: F_2,137_ = 5.73, *p*=0.0041], indicating greater analgesia after vapor exposure compared to CAN150 (post-hoc: *p*=0.0048) and vehicle vapor-exposed mice (post-hoc: *p=*0.0239; Supplemental Figure S3D). For locomotor activity in the open field (Supplemental Figure S3F), there was a significant Age x Sex interaction [F_1,142_ = 4.25, *p*=0.0410]; however, none of the Tukey’s post-hoc comparisons were significant.

## Supplemental Tables

**Table S1.**
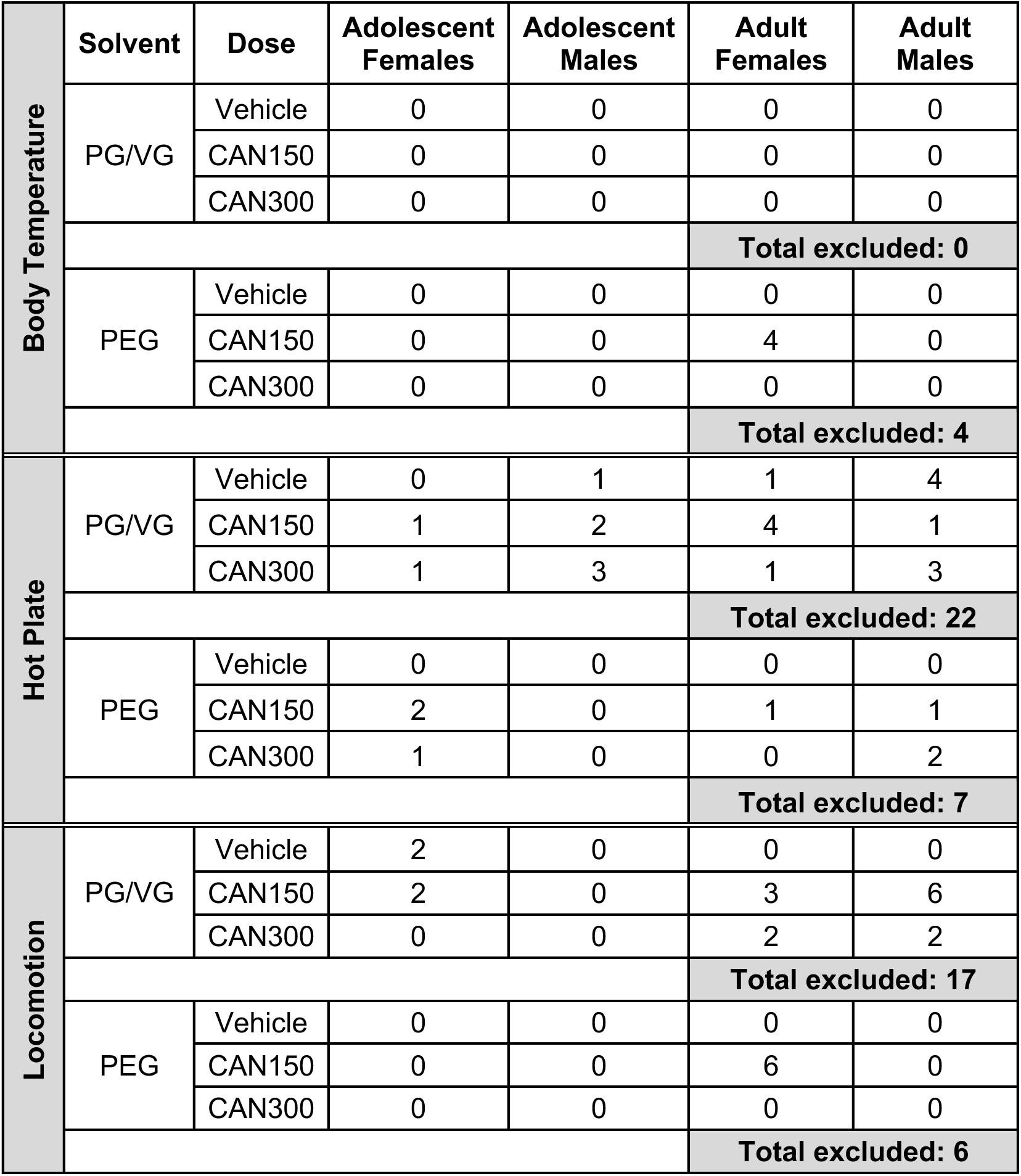
Group information for mice who met exclusion criteria for behavioral endpoints.

**Table S2.**
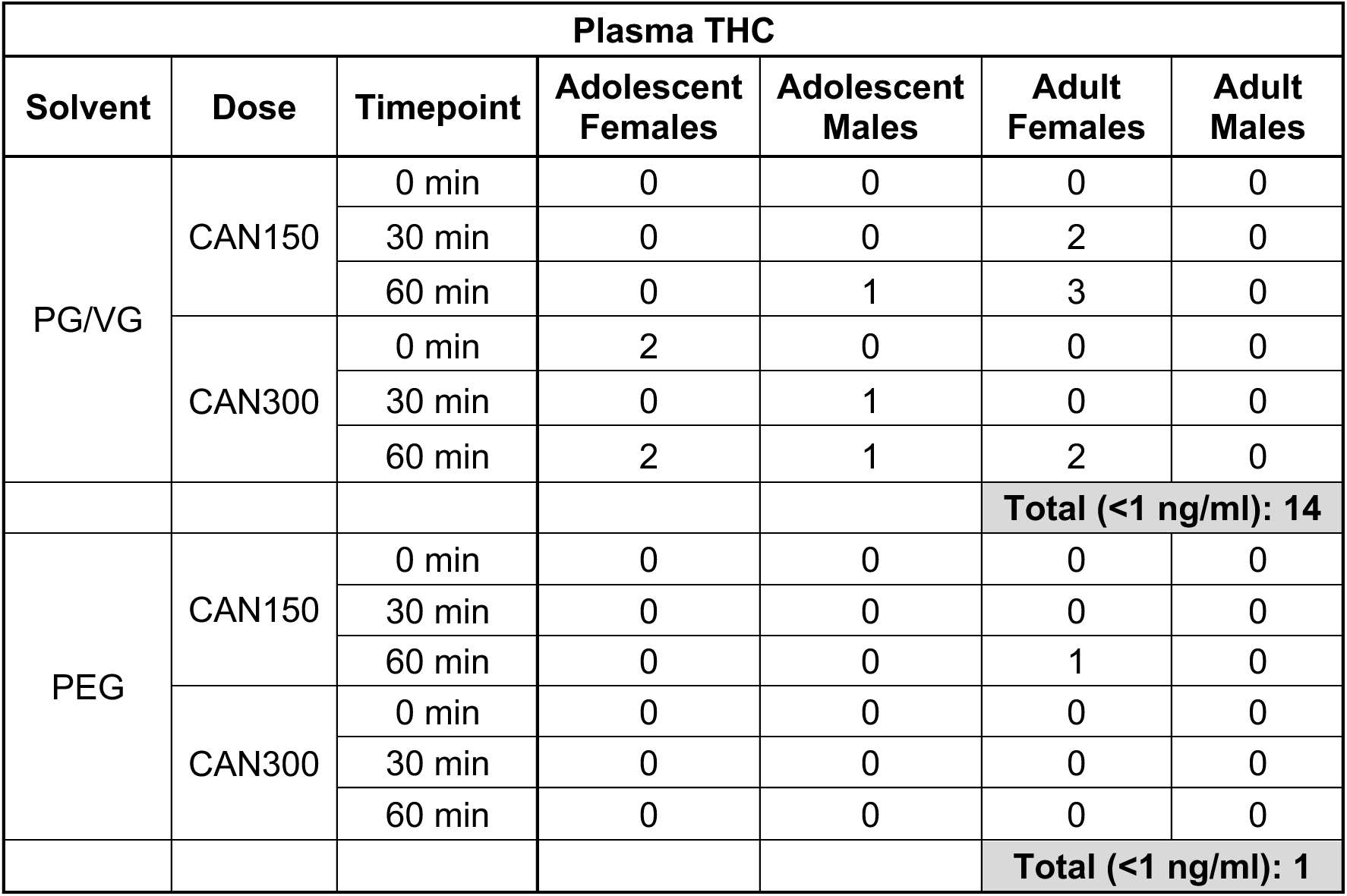
Number of plasma THC samples below the detection limit (<1 ng/ml) per group.

**Table S3.**
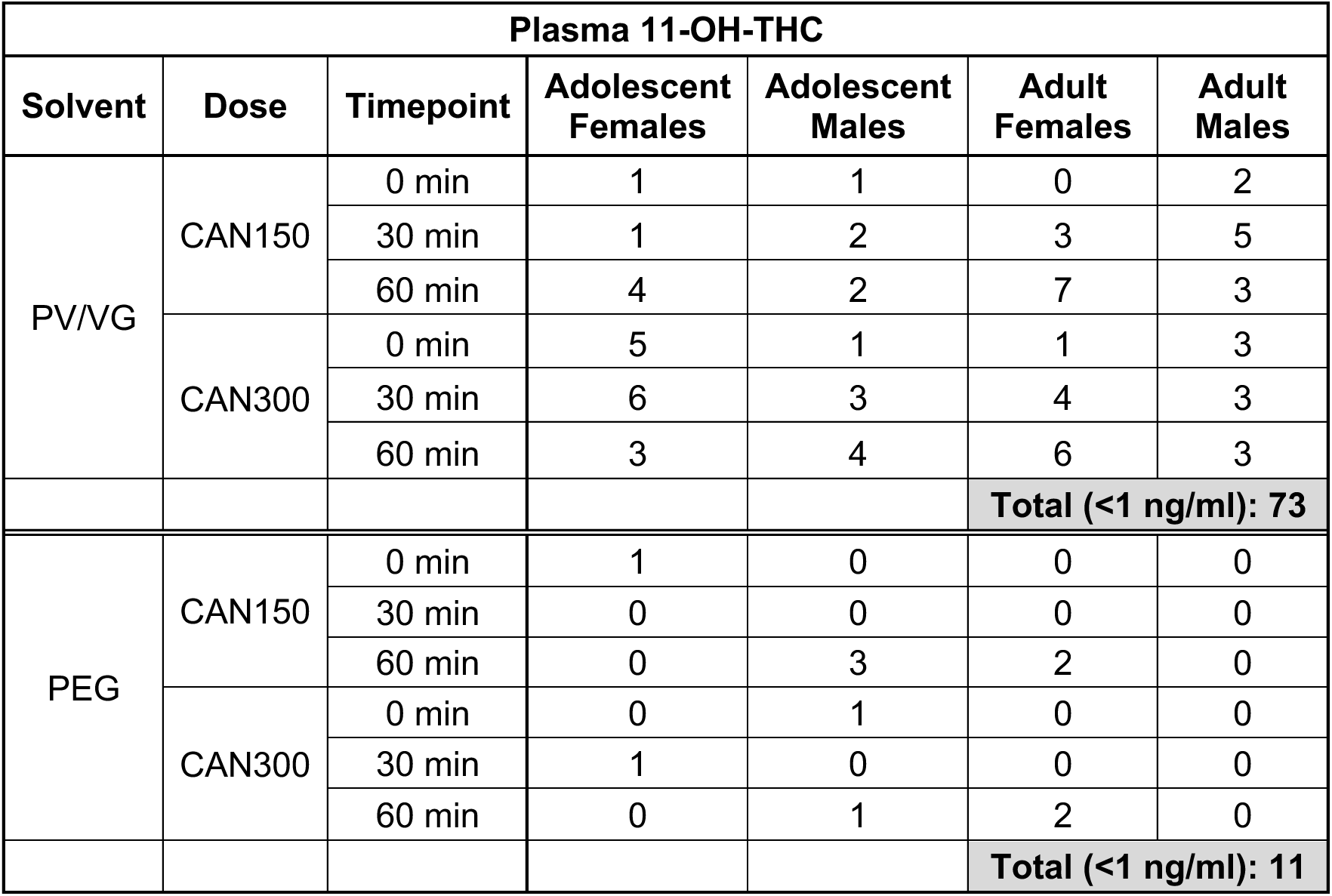
Number of plasma 11-OH-THC samples below the detection limit (<1 ng/ml) per group.

**Table S4.**
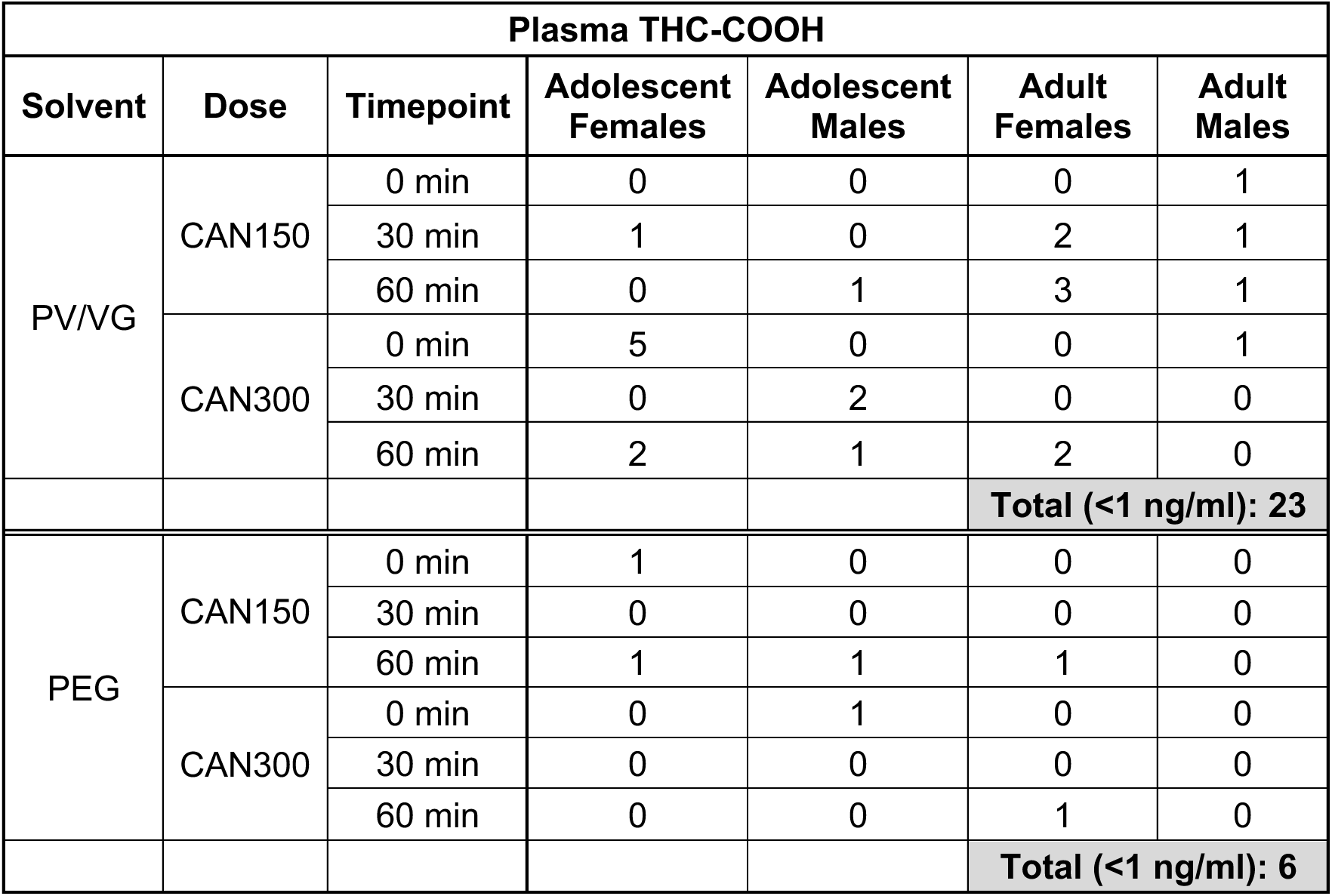
Number of plasma THC-COOH samples below the detection limit (<1 ng/ml) per group.

**Table S5.**
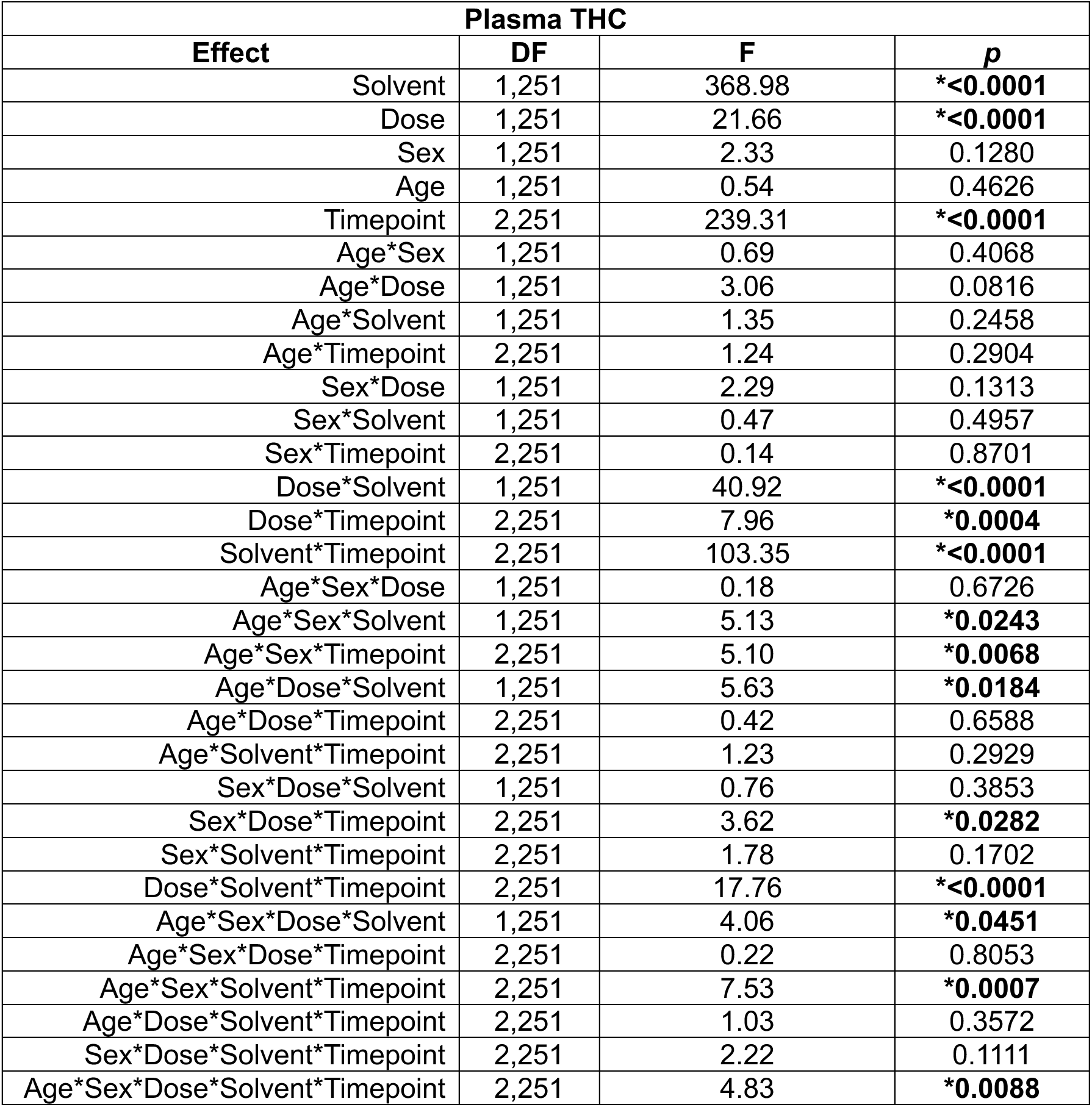
Statistical effects from a 5-way ANOVA with Solvent (PEG or PG/VG) as a factor for plasma THC levels.

**Table S6.**
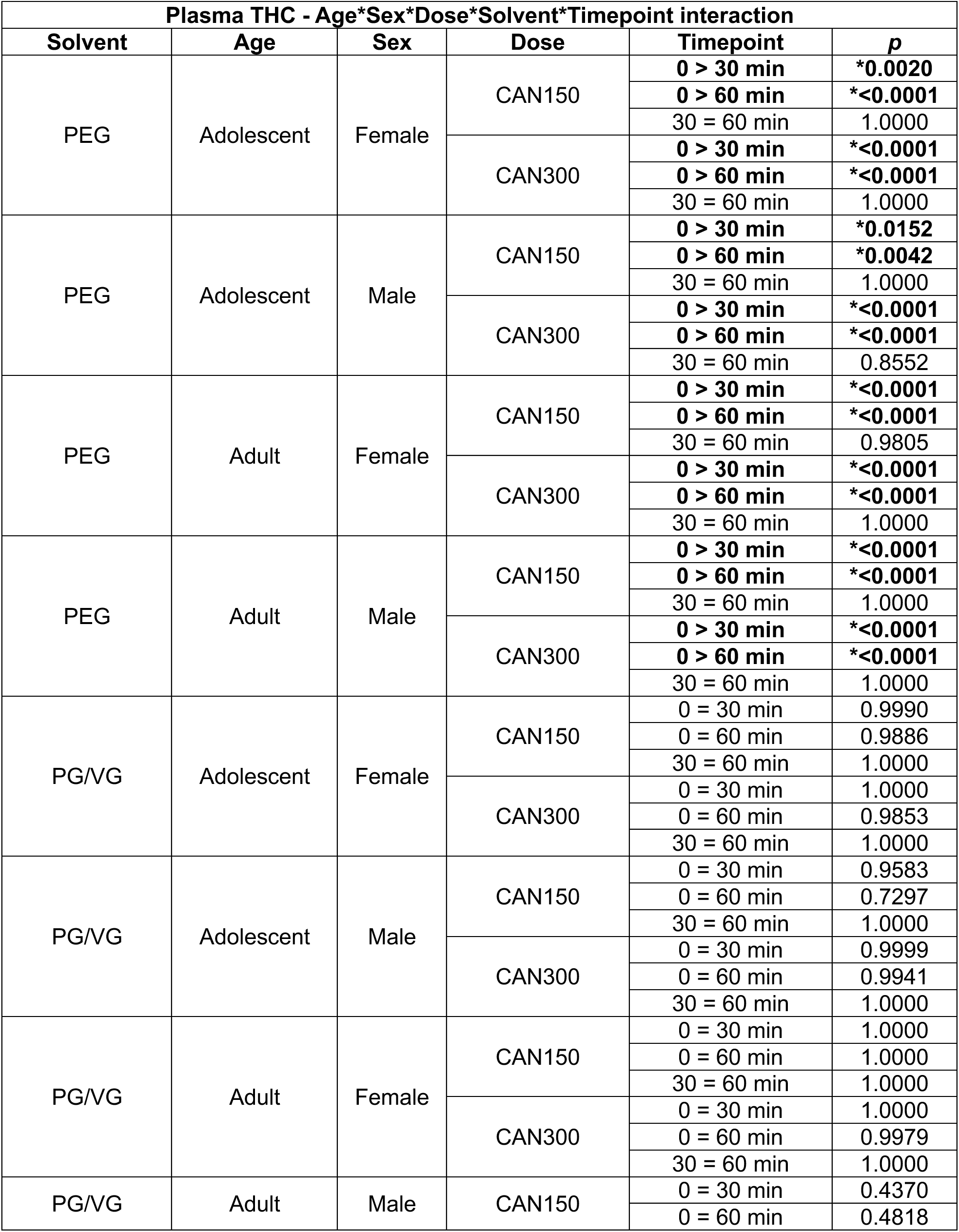
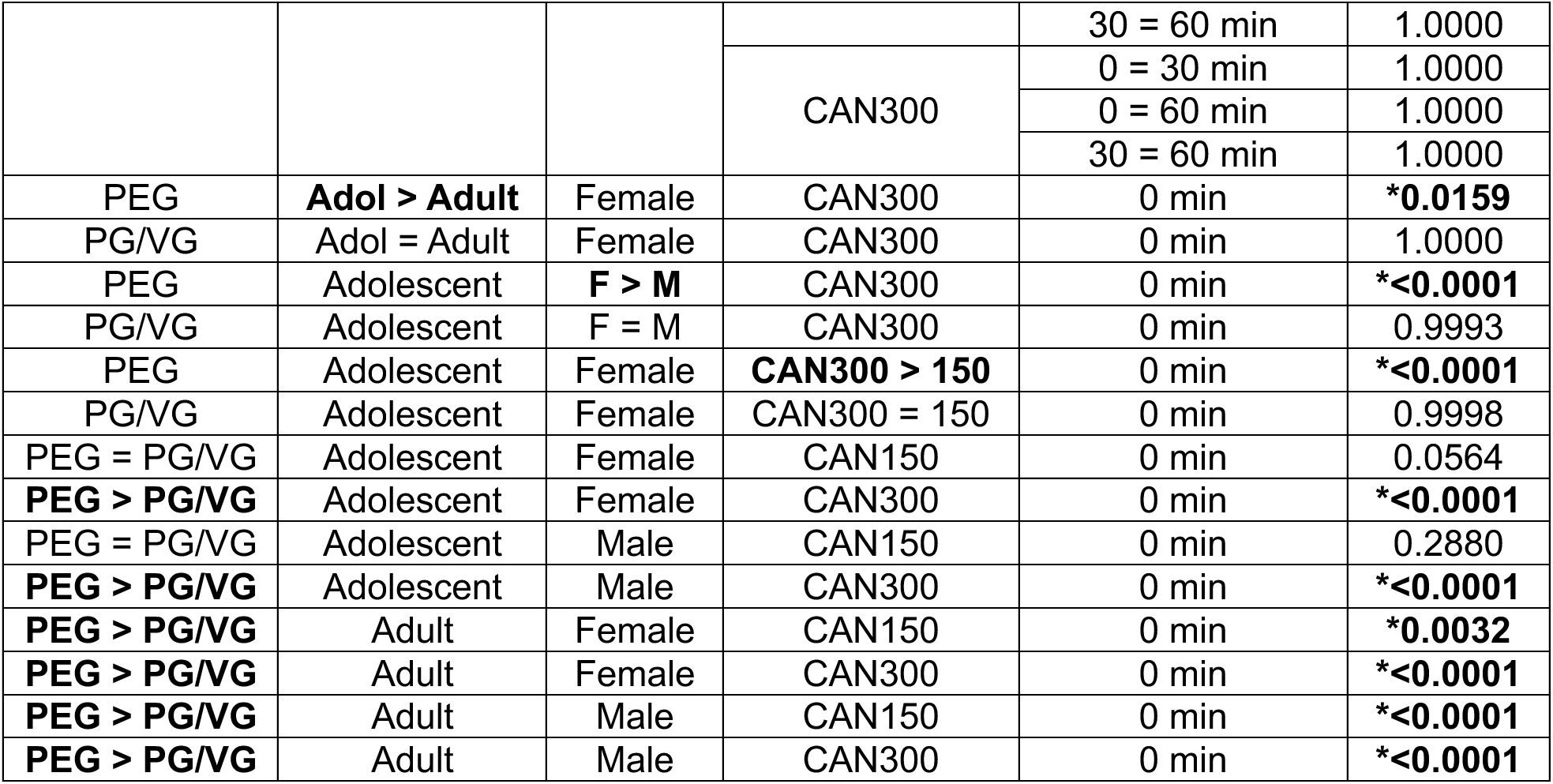
Notable Tukey’s post-hoc comparisons for plasma THC from the significant 5-way interaction in Table S5. Significant comparisons and their *p*-value are in bold.

**Table S7.**
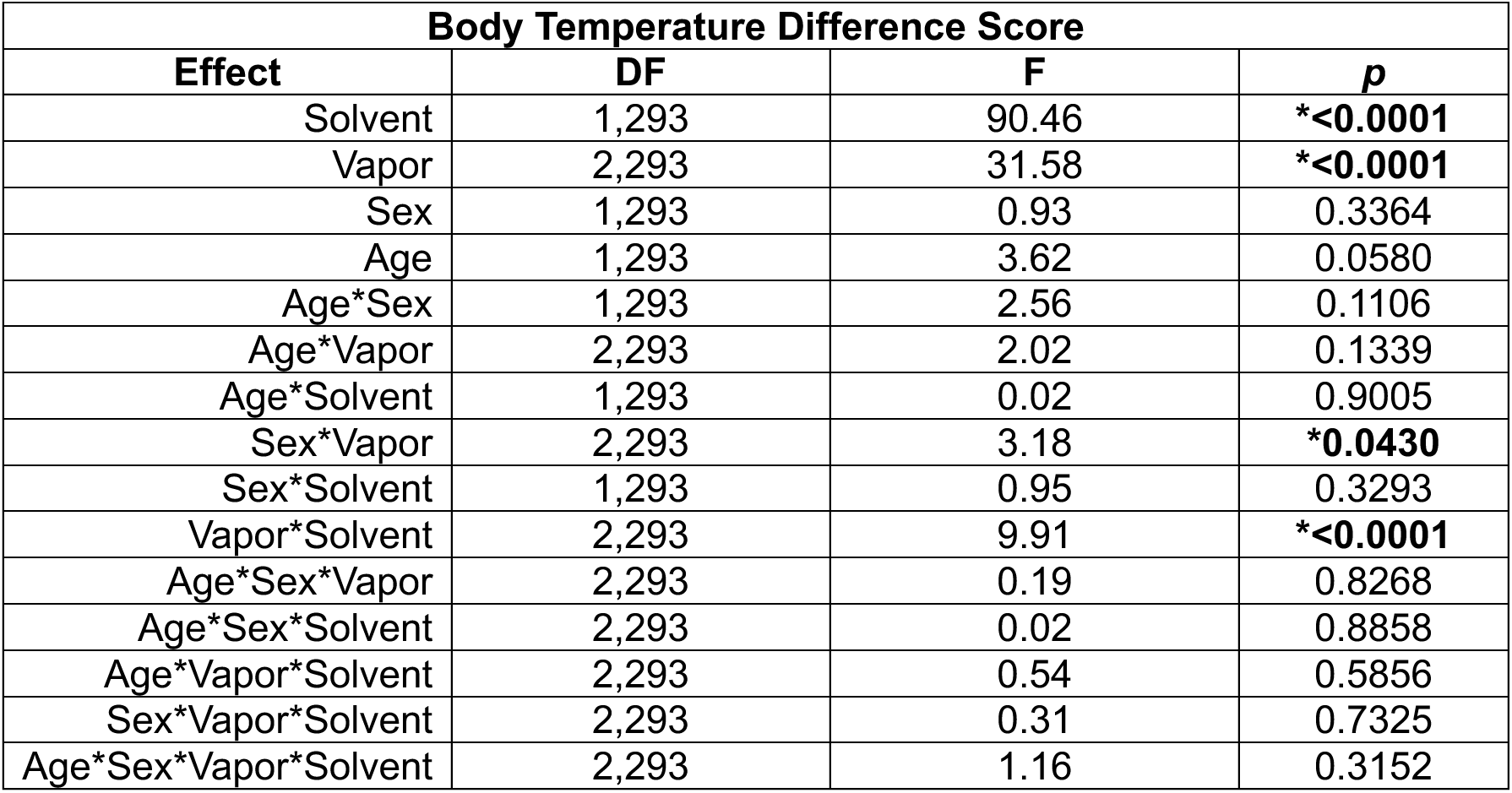
Statistical effects from a 4-way ANOVA with Solvent (PEG or PG/VG) as a factor for body temperature difference scores (post-exposure body temperature – pre-exposure body temperature).

**Table S8.**
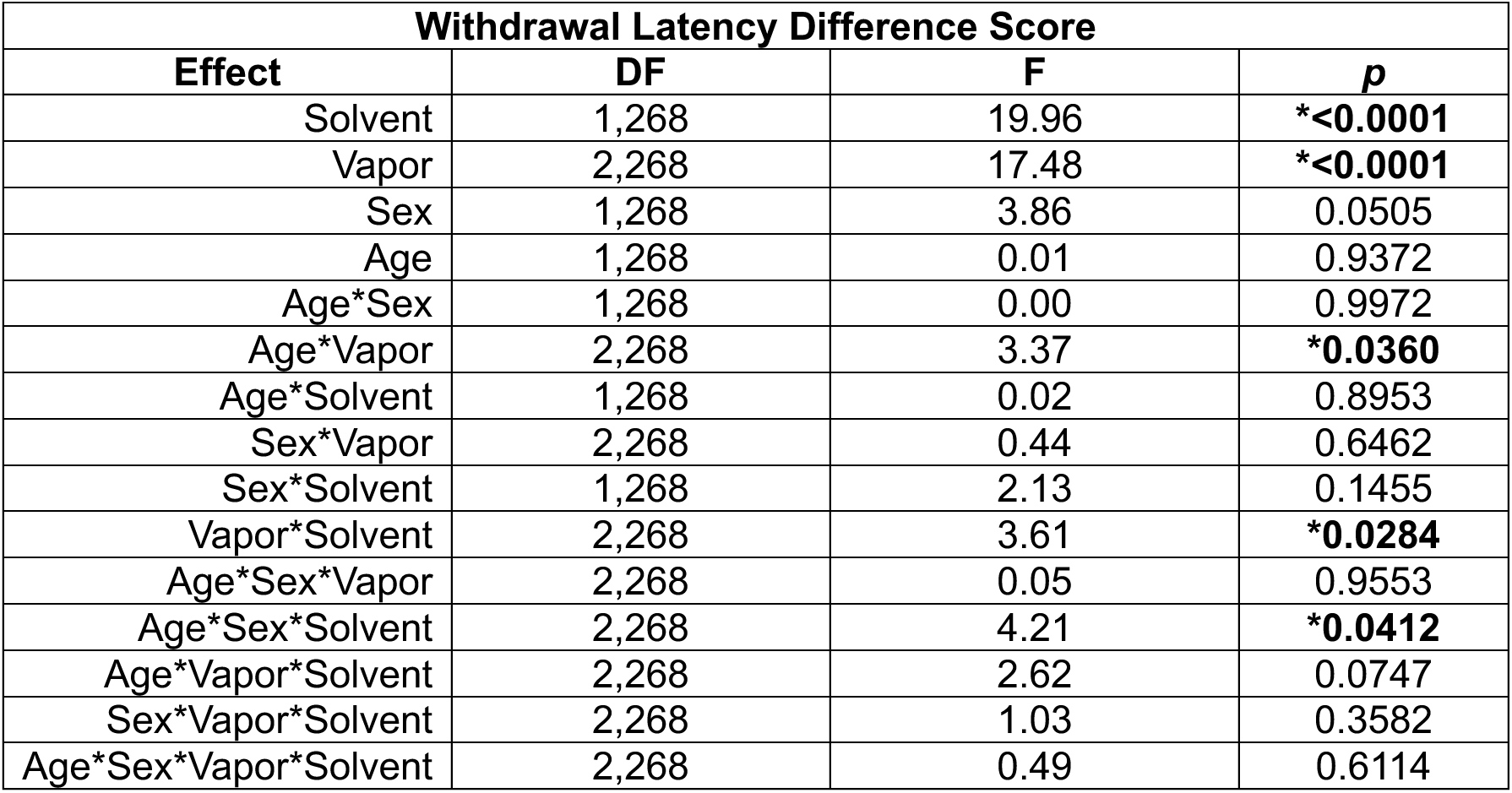
Statistical effects from a 4-way ANOVA with Solvent (PEG or PG/VG) as a factor for withdrawal latency difference scores (post-exposure latency – pre-exposure latency) in the hot plate test.

**Table S9.**
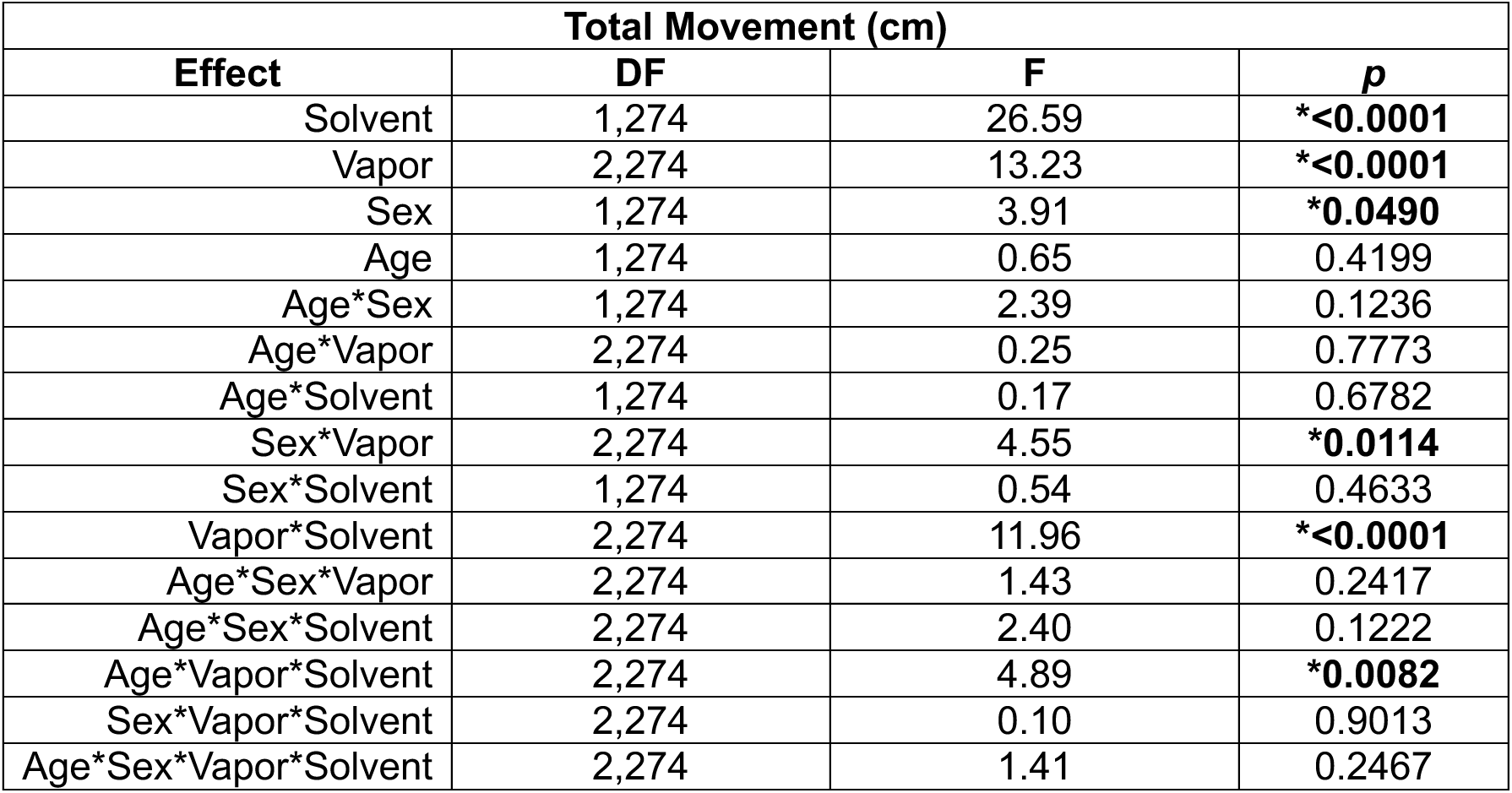
Statistical effects from a 4-way ANOVA with Solvent (PEG or PG/VG) as a factor for total movement (cm) in the open field.

## Supplemental Figures

**Figure S1.**
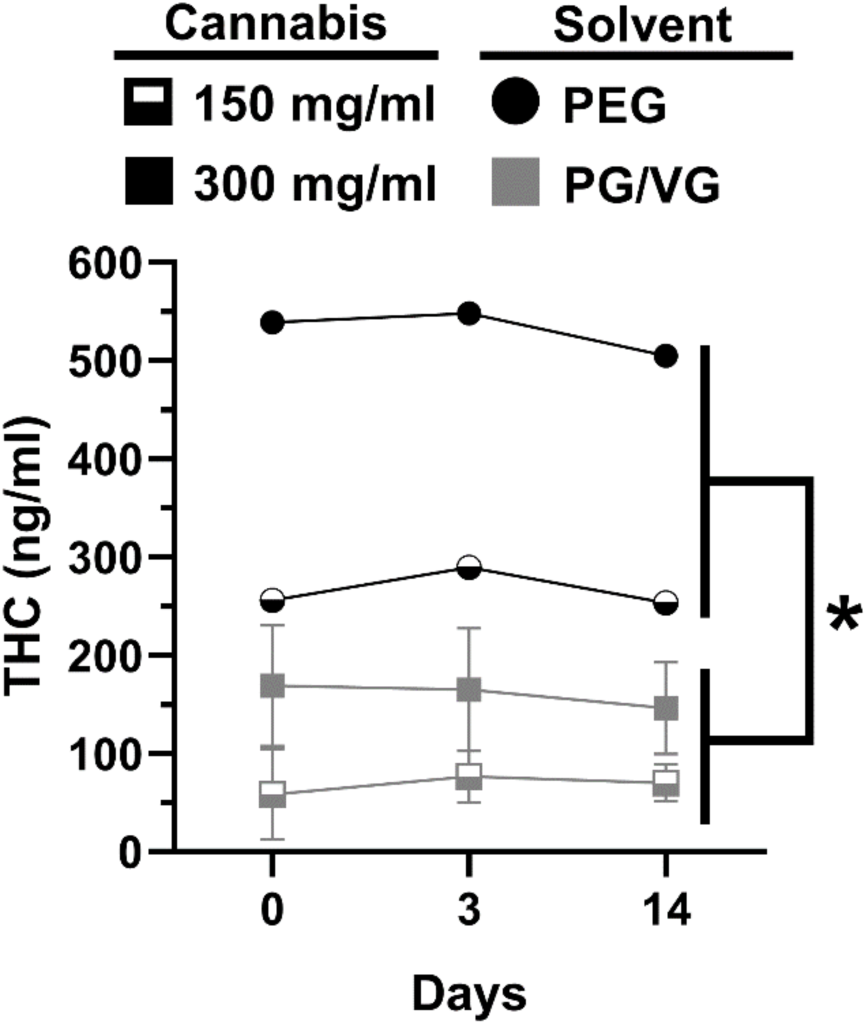
Time course of THC levels (ng/ml) in cannabis extract solutions (150 mg/ml, 300 mg/ml) made with different solvents (PEG, PG/VG). THC levels were significantly higher in cannabis extract solutions dissolved in PEG compared to PG/VG (main effect of Solvent: \**p*<0.05).

**Figure S2.**
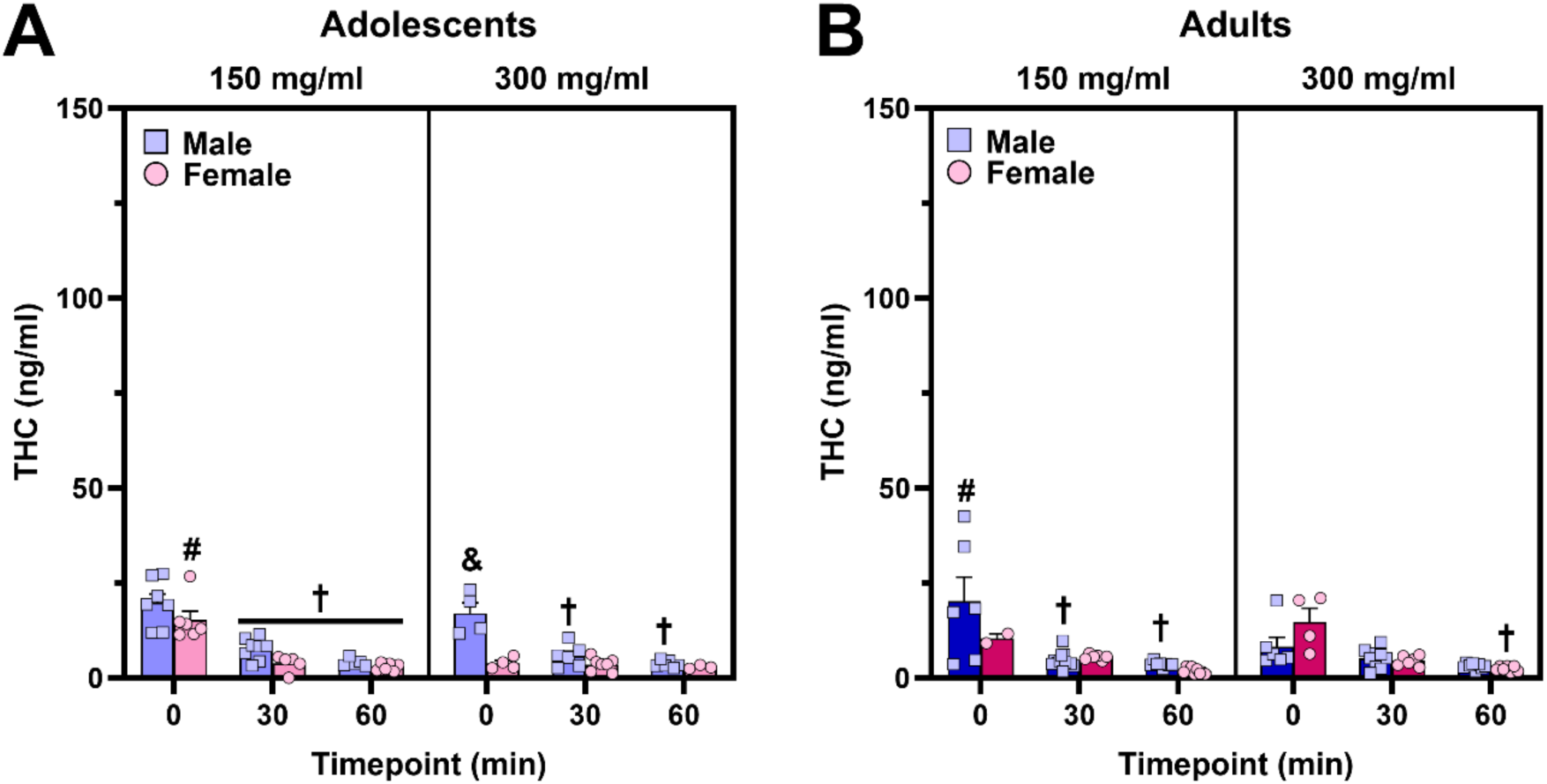
Time course of plasma THC and metabolites following 30-min vapor exposure session with PG/VG as the solvent. (**A,B**) Plasma THC levels when mice were exposed to cannabis dissolved in PG/VG showed time-dependent decreases (*post-hoc*: ^†^p<0.05, vs. 0 min timepoint within age, sex, and dose), although this was not present in all groups at both doses. There were unexpected dose-dependent *decreases* in plasma THC when comparing CAN150 and CAN300 groups (*post-hoc*: ^#^p<0.05, vs. CAN300 within sex, age, and timepoint). Adolescent male mice exposed to 300 mg/ml cannabis vapor reached higher plasma THC levels relative to their female counterparts at the 0 min timepoint (*post-hoc*: ^&^p<0.01, vs. adolescent females at the 0 min timepoint).

**Figure S3.**
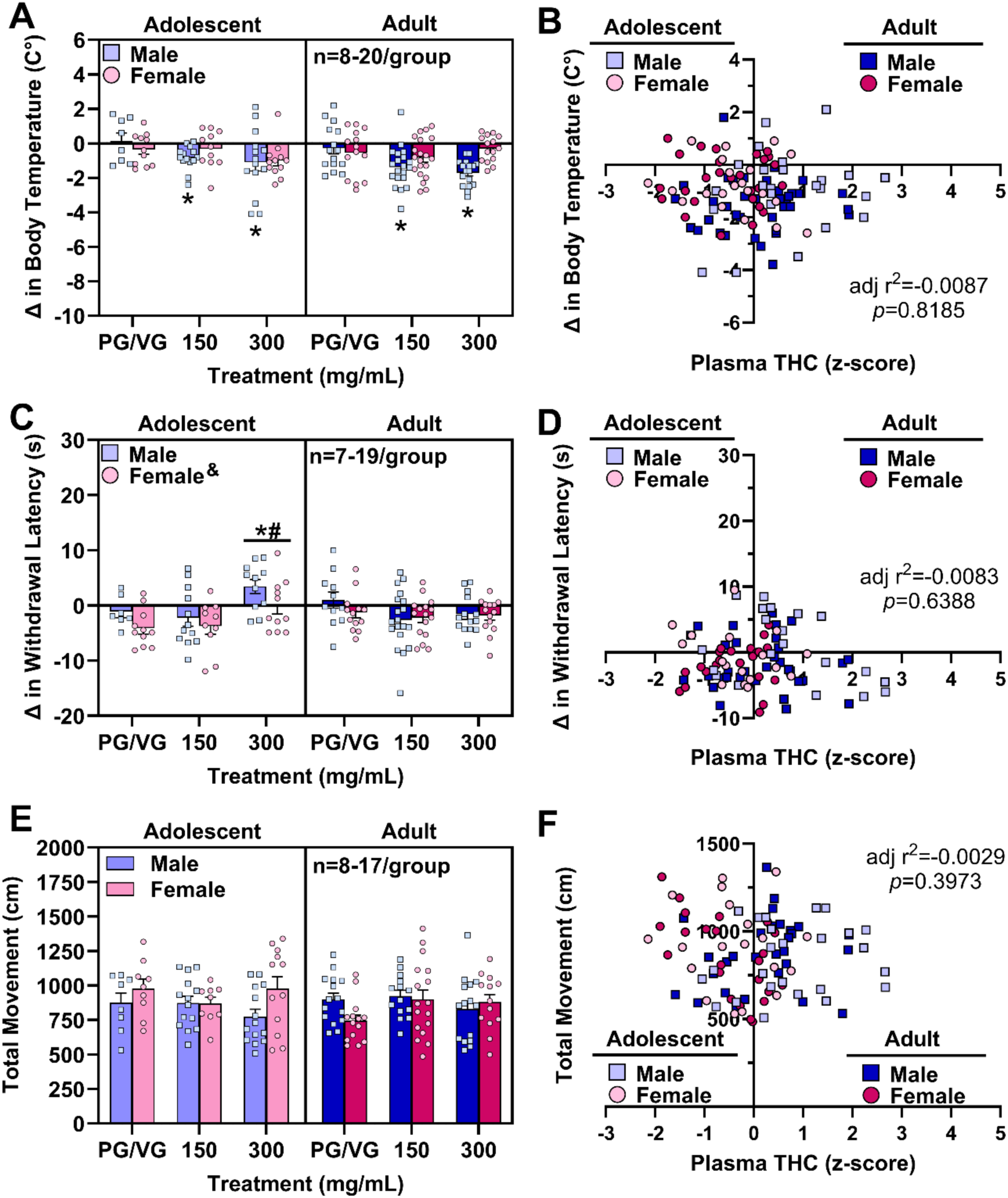
Vaporized cannabis extract dissolved in PG/VG induced hypothermia in male mice and antinociception in adolescent mice; however, plasma THC levels were not significantly correlated with behavioral endpoints. (**A**) Male mice exposed to 150 and 300 mg/ml cannabis vapor exhibited hypothermia relative to mice exposed to vehicle vapor (*post-hoc:* \**p’s*<0.05), regardless of age. (**B**) Plasma THC levels did not significantly predict changes in body temperature. (**C**) Adolescent mice exposed to 300 mg/ml cannabis vapor displayed greater antinociception relative to mice exposed to the lower dose of cannabis vapor (*post-hoc:* ^#^*p*<0.01) or vehicle vapor (*post-hoc:* \**p*<0.05), regardless of sex. Males had higher withdrawal latency difference scores than females (main effect of Sex: ^&^*p*<0.05). (**D**) Plasma THC levels did not significantly predict changes in withdrawal latency in the hot plate test. (**E**) Cannabis vapor at either dose did not significantly impact locomotor activity. (**F**) Plasma THC levels did not significantly predict locomotor activity.

## Notes

### Competing Interest Statement

The authors have declared no competing interest.

