## Supplemental results for "Influence of solvent, sex, and age on pharmacokinetic and acute behavioral effects of vaporized cannabis extract in mice"

**Supplemental Tables**

**Table S1.** Group information for mice who met exclusion criteria for behavioral endpoints.

| | Solvent | Dose | Adolescent Females | Adolescent Males | Adult Females | Adult Males |
|---|---|---|---|---|---|---|
| **Body Temperature** | PG/VG | Vehicle | 0 | 0 | 0 | 0 |
| | | CAN150 | 0 | 0 | 0 | 0 |
| | | CAN300 | 0 | 0 | 0 | 0 |
| | | | | | **Total excluded: 0** | |
| | PEG | Vehicle | 0 | 0 | 0 | 0 |
| | | CAN150 | 0 | 0 | 4 | 0 |
| | | CAN300 | 0 | 0 | 0 | 0 |
| | | | | | **Total excluded: 4** | |
| **Hot Plate** | PG/VG | Vehicle | 0 | 1 | 1 | 4 |
| | | CAN150 | 1 | 2 | 4 | 1 |
| | | CAN300 | 1 | 3 | 1 | 3 |
| | | | | | **Total excluded: 22** | |
| | PEG | Vehicle | 0 | 0 | 0 | 0 |
| | | CAN150 | 2 | 0 | 1 | 1 |
| | | CAN300 | 1 | 0 | 0 | 2 |
| | | | | | **Total excluded: 7** | |
| **Locomotion** | PG/VG | Vehicle | 2 | 0 | 0 | 0 |
| | | CAN150 | 2 | 0 | 3 | 6 |
| | | CAN300 | 0 | 0 | 2 | 2 |
| | | | | | **Total excluded: 17** | |
| | PEG | Vehicle | 0 | 0 | 0 | 0 |
| | | CAN150 | 0 | 0 | 6 | 0 |
| | | CAN300 | 0 | 0 | 0 | 0 |
| | | | | | **Total excluded: 6** | |

**Table S2.** Number of plasma THC samples below the detection limit (<1 ng/ml) per group.

| Plasma THC | | | | | | |
|---|---|---|---|---|---|---|
| **Solvent** | **Dose** | **Timepoint** | **Adolescent Females** | **Adolescent Males** | **Adult Females** | **Adult Males** |
| PG/VG | CAN150 | 0 min | 0 | 0 | 0 | 0 |
| | | 30 min | 0 | 0 | 2 | 0 |
| | | 60 min | 0 | 1 | 3 | 0 |
| | CAN300 | 0 min | 2 | 0 | 0 | 0 |
| | | 30 min | 0 | 1 | 0 | 0 |
| | | 60 min | 2 | 1 | 2 | 0 |
| | | | | | **Total (<1 ng/ml): 14** | |
| PEG | CAN150 | 0 min | 0 | 0 | 0 | 0 |
| | | 30 min | 0 | 0 | 0 | 0 |
| | | 60 min | 0 | 0 | 1 | 0 |
| | CAN300 | 0 min | 0 | 0 | 0 | 0 |
| | | 30 min | 0 | 0 | 0 | 0 |
| | | 60 min | 0 | 0 | 0 | 0 |
| | | | | | **Total (<1 ng/ml): 1** | |

**Table S3.** Number of plasma 11-OH-THC samples below the detection limit (<1 ng/ml) per group.

| Plasma 11-OH-THC | | | | | | |
|---|---|---|---|---|---|---|
| **Solvent** | **Dose** | **Timepoint** | **Adolescent Females** | **Adolescent Males** | **Adult Females** | **Adult Males** |
| PV/VG | CAN150 | 0 min | 1 | 1 | 0 | 2 |
| | | 30 min | 1 | 2 | 3 | 5 |
| | | 60 min | 4 | 2 | 7 | 3 |
| | CAN300 | 0 min | 5 | 1 | 1 | 3 |
| | | 30 min | 6 | 3 | 4 | 3 |
| | | 60 min | 3 | 4 | 6 | 3 |
| | | | | | **Total (<1 ng/ml): 73** | |
| PEG | CAN150 | 0 min | 1 | 0 | 0 | 0 |
| | | 30 min | 0 | 0 | 0 | 0 |
| | | 60 min | 0 | 3 | 2 | 0 |
| | CAN300 | 0 min | 0 | 1 | 0 | 0 |
| | | 30 min | 1 | 0 | 0 | 0 |
| | | 60 min | 0 | 1 | 2 | 0 |
| | | | | | **Total (<1 ng/ml): 11** | |

**Table S4.** Number of plasma THC-COOH samples below the detection limit (<1 ng/ml) per group.

| Plasma THC-COOH | | | | | | |
|---|---|---|---|---|---|---|
| **Solvent** | **Dose** | **Timepoint** | **Adolescent Females** | **Adolescent Males** | **Adult Females** | **Adult Males** |
| PV/VG | CAN150 | 0 min | 0 | 0 | 0 | 1 |
| | | 30 min | 1 | 0 | 2 | 1 |
| | | 60 min | 0 | 1 | 3 | 1 |
| | CAN300 | 0 min | 5 | 0 | 0 | 1 |
| | | 30 min | 0 | 2 | 0 | 0 |
| | | 60 min | 2 | 1 | 2 | 0 |
| | | | | | **Total (<1 ng/ml): 23** | |
| PEG | CAN150 | 0 min | 1 | 0 | 0 | 0 |
| | | 30 min | 0 | 0 | 0 | 0 |
| | | 60 min | 1 | 1 | 1 | 0 |
| | CAN300 | 0 min | 0 | 1 | 0 | 0 |
| | | 30 min | 0 | 0 | 0 | 0 |
| | | 60 min | 0 | 0 | 1 | 0 |
| | | | | | **Total (<1 ng/ml): 6** | |

**Table S5.** Statistical effects from a 5-way ANOVA with Solvent (PEG or PG/VG) as a factor for plasma THC levels.

| Plasma THC | | | |
|---|---|---|---|
| **Effect** | **DF** | **F** | ***p*** |
| Solvent | 1,251 | 368.98 | ***<0.0001** |
| Dose | 1,251 | 21.66 | ***<0.0001** |
| Sex | 1,251 | 2.33 | 0.1280 |
| Age | 1,251 | 0.54 | 0.4626 |
| Timepoint | 2,251 | 239.31 | ***<0.0001** |
| Age*Sex | 1,251 | 0.69 | 0.4068 |
| Age*Dose | 1,251 | 3.06 | 0.0816 |
| Age*Solvent | 1,251 | 1.35 | 0.2458 |
| Age*Timepoint | 2,251 | 1.24 | 0.2904 |
| Sex*Dose | 1,251 | 2.29 | 0.1313 |
| Sex*Solvent | 1,251 | 0.47 | 0.4957 |
| Sex*Timepoint | 2,251 | 0.14 | 0.8701 |
| Dose*Solvent | 1,251 | 40.92 | ***<0.0001** |
| Dose*Timepoint | 2,251 | 7.96 | ***0.0004** |
| Solvent*Timepoint | 2,251 | 103.35 | ***<0.0001** |
| Age*Sex*Dose | 1,251 | 0.18 | 0.6726 |
| Age*Sex*Solvent | 1,251 | 5.13 | ***0.0243** |
| Age*Sex*Timepoint | 2,251 | 5.10 | ***0.0068** |
| Age*Dose*Solvent | 1,251 | 5.63 | ***0.0184** |
| Age*Dose*Timepoint | 2,251 | 0.42 | 0.6588 |
| Age*Solvent*Timepoint | 2,251 | 1.23 | 0.2929 |
| Sex*Dose*Solvent | 1,251 | 0.76 | 0.3853 |
| Sex*Dose*Timepoint | 2,251 | 3.62 | ***0.0282** |
| Sex*Solvent*Timepoint | 2,251 | 1.78 | 0.1702 |
| Dose*Solvent*Timepoint | 2,251 | 17.76 | ***<0.0001** |
| Age*Sex*Dose*Solvent | 1,251 | 4.06 | ***0.0451** |
| Age*Sex*Dose*Timepoint | 2,251 | 0.22 | 0.8053 |
| Age*Sex*Solvent*Timepoint | 2,251 | 7.53 | ***0.0007** |
| Age*Dose*Solvent*Timepoint | 2,251 | 1.03 | 0.3572 |
| Sex*Dose*Solvent*Timepoint | 2,251 | 2.22 | 0.1111 |
| Age*Sex*Dose*Solvent*Timepoint | 2,251 | 4.83 | ***0.0088** |

**Table S6.** Notable Tukey's post-hoc comparisons for plasma THC from the significant 5-way interaction in Table S5. Significant comparisons and their *p*-value are in bold.

| **Plasma THC - Age*Sex*Dose*Solvent*Timepoint interaction** | | | | | |
|---|---|---|---|---|---|
| **Solvent** | **Age** | **Sex** | **Dose** | **Timepoint** | ***p*** |
| PEG | Adolescent | Female | CAN150 | **0 > 30 min** | ***0.0020** |
| | | | | **0 > 60 min** | ***<0.0001** |
| | | | | 30 = 60 min | 1.0000 |
| | | | CAN300 | **0 > 30 min** | ***<0.0001** |
| | | | | **0 > 60 min** | ***<0.0001** |
| | | | | 30 = 60 min | 1.0000 |
| PEG | Adolescent | Male | CAN150 | **0 > 30 min** | ***0.0152** |
| | | | | **0 > 60 min** | ***0.0042** |
| | | | | 30 = 60 min | 1.0000 |
| | | | CAN300 | **0 > 30 min** | ***<0.0001** |
| | | | | **0 > 60 min** | ***<0.0001** |
| | | | | 30 = 60 min | 0.8552 |
| PEG | Adult | Female | CAN150 | **0 > 30 min** | ***<0.0001** |
| | | | | **0 > 60 min** | ***<0.0001** |
| | | | | 30 = 60 min | 0.9805 |
| | | | CAN300 | **0 > 30 min** | ***<0.0001** |
| | | | | **0 > 60 min** | ***<0.0001** |
| | | | | 30 = 60 min | 1.0000 |
| PEG | Adult | Male | CAN150 | **0 > 30 min** | ***<0.0001** |
| | | | | **0 > 60 min** | ***<0.0001** |
| | | | | 30 = 60 min | 1.0000 |
| | | | CAN300 | **0 > 30 min** | ***<0.0001** |
| | | | | **0 > 60 min** | ***<0.0001** |
| | | | | 30 = 60 min | 1.0000 |
| PG/VG | Adolescent | Female | CAN150 | 0 = 30 min | 0.9990 |
| | | | | 0 = 60 min | 0.9886 |
| | | | | 30 = 60 min | 1.0000 |
| | | | CAN300 | 0 = 30 min | 1.0000 |
| | | | | 0 = 60 min | 0.9853 |
| | | | | 30 = 60 min | 1.0000 |
| PG/VG | Adolescent | Male | CAN150 | 0 = 30 min | 0.9583 |
| | | | | 0 = 60 min | 0.7297 |
| | | | | 30 = 60 min | 1.0000 |
| | | | CAN300 | 0 = 30 min | 0.9999 |
| | | | | 0 = 60 min | 0.9941 |
| | | | | 30 = 60 min | 1.0000 |
| PG/VG | Adult | Female | CAN150 | 0 = 30 min | 1.0000 |
| | | | | 0 = 60 min | 1.0000 |
| | | | | 30 = 60 min | 1.0000 |
| | | | CAN300 | 0 = 30 min | 1.0000 |
| | | | | 0 = 60 min | 0.9979 |
| | | | | 30 = 60 min | 1.0000 |
| PG/VG | Adult | Male | CAN150 | 0 = 30 min | 0.4370 |
| | | | | 0 = 60 min | 0.4818 |

| | | | | | |
|---|---|---|---|---|---|
| | | | | 30 = 60 min | 1.0000 |
| | | | CAN300 | 0 = 30 min | 1.0000 |
| | | | | 0 = 60 min | 1.0000 |
| | | | | 30 = 60 min | 1.0000 |
| PEG | **Adol > Adult** | Female | CAN300 | 0 min | ***0.0159** |
| PG/VG | Adol = Adult | Female | CAN300 | 0 min | 1.0000 |
| PEG | Adolescent | **F > M** | CAN300 | 0 min | ***<0.0001** |
| PG/VG | Adolescent | F = M | CAN300 | 0 min | 0.9993 |
| PEG | Adolescent | Female | **CAN300 > 150** | 0 min | ***<0.0001** |
| PG/VG | Adolescent | Female | CAN300 = 150 | 0 min | 0.9998 |
| PEG = PG/VG | Adolescent | Female | CAN150 | 0 min | 0.0564 |
| **PEG > PG/VG** | Adolescent | Female | CAN300 | 0 min | ***<0.0001** |
| PEG = PG/VG | Adolescent | Male | CAN150 | 0 min | 0.2880 |
| **PEG > PG/VG** | Adolescent | Male | CAN300 | 0 min | ***<0.0001** |
| **PEG > PG/VG** | Adult | Female | CAN150 | 0 min | ***0.0032** |
| **PEG > PG/VG** | Adult | Female | CAN300 | 0 min | ***<0.0001** |
| **PEG > PG/VG** | Adult | Male | CAN150 | 0 min | ***<0.0001** |
| **PEG > PG/VG** | Adult | Male | CAN300 | 0 min | ***<0.0001** |

**Table S7.** Statistical effects from a 4-way ANOVA with Solvent (PEG or PG/VG) as a factor for body temperature difference scores (post-exposure body temperature – pre-exposure body temperature).

| **Body Temperature Difference Score** | | | |
|---|---|---|---|
| **Effect** | **DF** | **F** | ***p*** |
| Solvent | 1,293 | 90.46 | ***<0.0001** |
| Vapor | 2,293 | 31.58 | ***<0.0001** |
| Sex | 1,293 | 0.93 | 0.3364 |
| Age | 1,293 | 3.62 | 0.0580 |
| Age*Sex | 1,293 | 2.56 | 0.1106 |
| Age*Vapor | 2,293 | 2.02 | 0.1339 |
| Age*Solvent | 1,293 | 0.02 | 0.9005 |
| Sex*Vapor | 2,293 | 3.18 | ***0.0430** |
| Sex*Solvent | 1,293 | 0.95 | 0.3293 |
| Vapor*Solvent | 2,293 | 9.91 | ***<0.0001** |
| Age*Sex*Vapor | 2,293 | 0.19 | 0.8268 |
| Age*Sex*Solvent | 2,293 | 0.02 | 0.8858 |
| Age*Vapor*Solvent | 2,293 | 0.54 | 0.5856 |
| Sex*Vapor*Solvent | 2,293 | 0.31 | 0.7325 |
| Age*Sex*Vapor*Solvent | 2,293 | 1.16 | 0.3152 |

**Table S8.** Statistical effects from a 4-way ANOVA with Solvent (PEG or PG/VG) as a factor for withdrawal latency difference scores (post-exposure latency – pre-exposure latency) in the hot plate test.

| **Withdrawal Latency Difference Score** | | | |
|---|---|---|---|
| **Effect** | **DF** | **F** | ***p*** |
| Solvent | 1,268 | 19.96 | ***<0.0001** |
| Vapor | 2,268 | 17.48 | ***<0.0001** |
| Sex | 1,268 | 3.86 | 0.0505 |
| Age | 1,268 | 0.01 | 0.9372 |
| Age*Sex | 1,268 | 0.00 | 0.9972 |
| Age*Vapor | 2,268 | 3.37 | ***0.0360** |
| Age*Solvent | 1,268 | 0.02 | 0.8953 |
| Sex*Vapor | 2,268 | 0.44 | 0.6462 |
| Sex*Solvent | 1,268 | 2.13 | 0.1455 |
| Vapor*Solvent | 2,268 | 3.61 | ***0.0284** |
| Age*Sex*Vapor | 2,268 | 0.05 | 0.9553 |
| Age*Sex*Solvent | 2,268 | 4.21 | ***0.0412** |
| Age*Vapor*Solvent | 2,268 | 2.62 | 0.0747 |
| Sex*Vapor*Solvent | 2,268 | 1.03 | 0.3582 |
| Age*Sex*Vapor*Solvent | 2,268 | 0.49 | 0.6114 |

**Table S9.** Statistical effects from a 4-way ANOVA with Solvent (PEG or PG/VG) as a factor for total movement (cm) in the open field.

| Total Movement (cm) | | | |
|---|---|---|---|
| **Effect** | **DF** | **F** | ***p*** |
| Solvent | 1,274 | 26.59 | ***<0.0001** |
| Vapor | 2,274 | 13.23 | ***<0.0001** |
| Sex | 1,274 | 3.91 | ***0.0490** |
| Age | 1,274 | 0.65 | 0.4199 |
| Age*Sex | 1,274 | 2.39 | 0.1236 |
| Age*Vapor | 2,274 | 0.25 | 0.7773 |
| Age*Solvent | 1,274 | 0.17 | 0.6782 |
| Sex*Vapor | 2,274 | 4.55 | ***0.0114** |
| Sex*Solvent | 1,274 | 0.54 | 0.4633 |
| Vapor*Solvent | 2,274 | 11.96 | ***<0.0001** |
| Age*Sex*Vapor | 2,274 | 1.43 | 0.2417 |
| Age*Sex*Solvent | 2,274 | 2.40 | 0.1222 |
| Age*Vapor*Solvent | 2,274 | 4.89 | ***0.0082** |
| Sex*Vapor*Solvent | 2,274 | 0.10 | 0.9013 |
| Age*Sex*Vapor*Solvent | 2,274 | 1.41 | 0.2467 |

**Supplemental Figures**

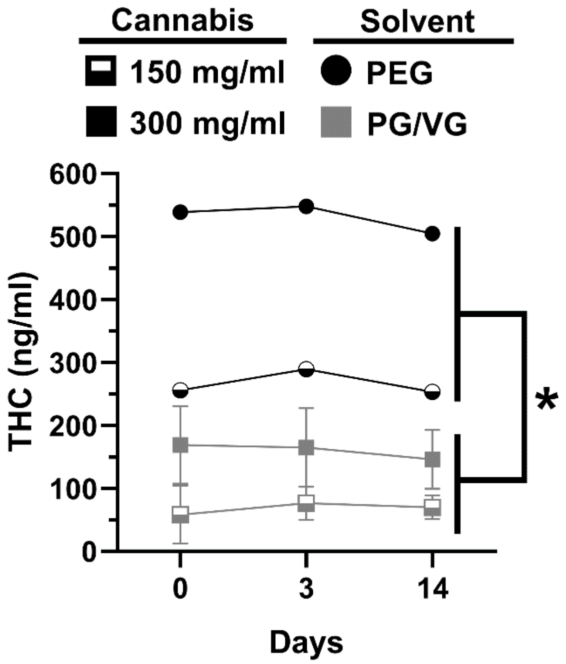


**Figure S1.** Time course of THC levels (ng/ml) in cannabis extract solutions (150 mg/ml, 300 mg/ml) made with different solvents (PEG, PG/VG). THC levels were significantly higher in cannabis extract solutions dissolved in PEG compared to PG/VG (main effect of Solvent: *$p<0.05$).

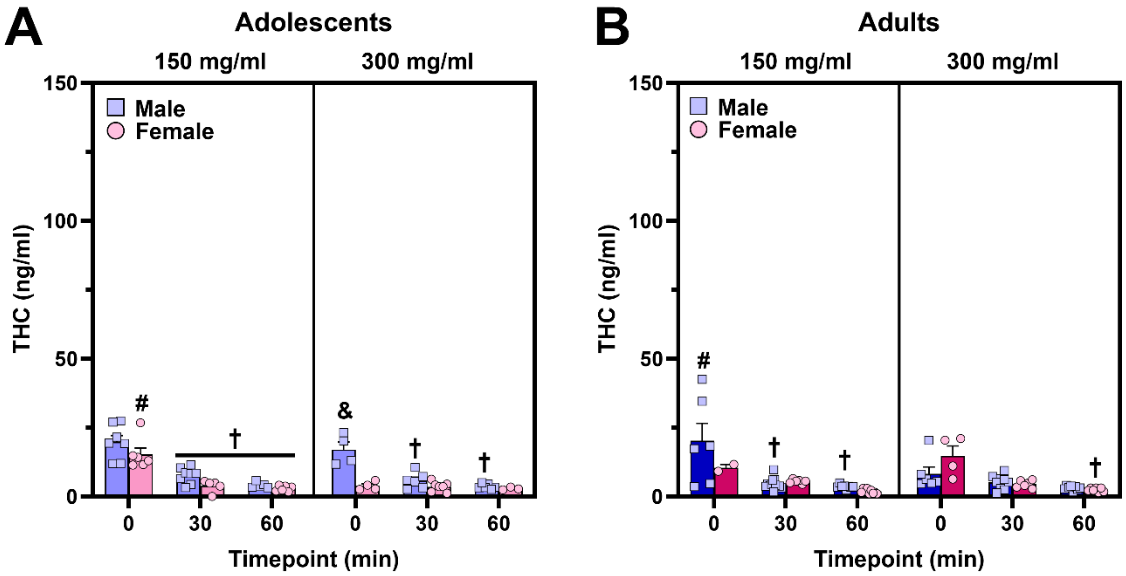


**Figure S2.** Time course of plasma THC and metabolites following 30-min vapor exposure session with PG/VG as the solvent. (**A,B**) Plasma THC levels when mice were exposed to cannabis dissolved in PG/VG showed time-dependent decreases (*post-hoc*: $^{\dagger}p<0.05$, vs. 0 min timepoint within age, sex, and dose), although this was not present in all groups at both doses. There were unexpected dose-dependent *decreases* in plasma THC when comparing CAN150 and CAN300 groups (*post-hoc*: $^{\#}p<0.05$, vs. CAN300 within sex, age, and timepoint). Adolescent male mice exposed to 300 mg/ml cannabis vapor reached higher plasma THC levels relative to their female counterparts at the 0 min timepoint (*post-hoc*: $^{\&}p<0.01$, vs. adolescent females at the 0 min timepoint).

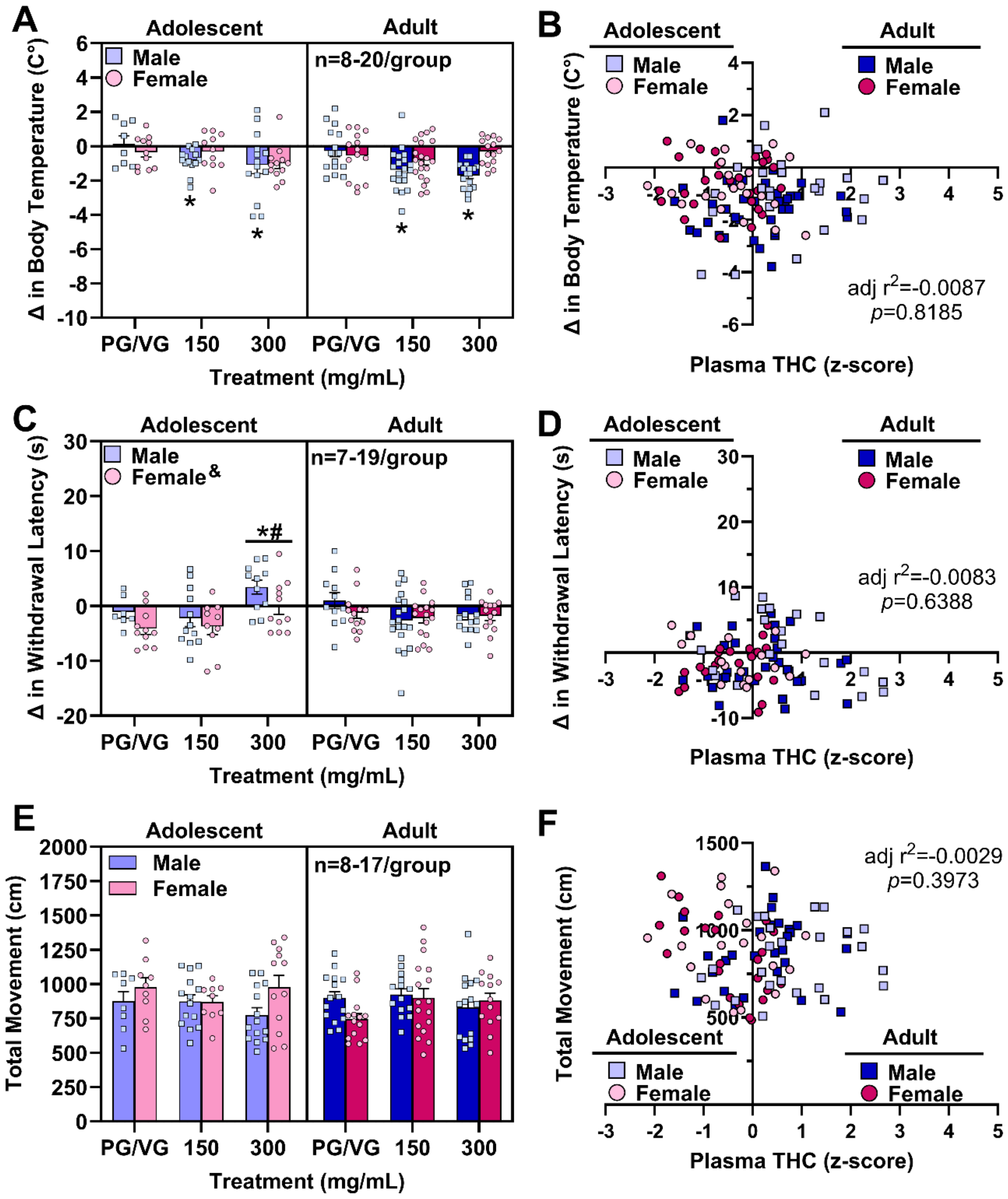


**Figure S3.** Vaporized cannabis extract dissolved in PG/VG induced hypothermia in male mice and antinociception in adolescent mice; however, plasma THC levels were not significantly correlated with behavioral endpoints. (**A**) Male mice exposed to 150 and 300 mg/ml cannabis vapor exhibited hypothermia relative to mice exposed to vehicle vapor (*post-hoc:* **p's*<0.05), regardless of age. (**B**) Plasma THC levels did not significantly predict changes in body temperature. (**C**) Adolescent mice exposed to 300 mg/ml cannabis vapor displayed greater antinociception relative to mice exposed to the lower dose of cannabis vapor (*post-hoc:* $^{\#}p<0.01$) or vehicle vapor (*post-hoc:* *$p<0.05$), regardless of sex. Males had higher withdrawal latency difference scores than females (main effect of Sex: $^{\&}p<0.05$). (**D**) Plasma THC levels did not significantly predict changes in withdrawal latency in the hot plate test. (**E**) Cannabis vapor at either dose did not significantly impact locomotor activity. (**F**) Plasma THC levels did not significantly predict locomotor activity.
